# Intracellular pocket conformations determine signaling efficacy through the μ opioid receptor

**DOI:** 10.1101/2024.04.03.588021

**Authors:** David A. Cooper, Joseph DePaolo-Boisvert, Stanley A. Nicholson, Barien Gad, David D. L. Minh

**Affiliations:** Department of Chemistry, Illinois Institute of Technology, Chicago, Illinois 60616, United States; Department of Applied Mathematics, Illinois Institute of Technology, Chicago, Illinois 60616, United States; Department of Biology, Illinois Institute of Technology, Chicago, Illinois 60616, United States

**Keywords:** 7 transmembrane helix receptor. G protein coupled receptor. Functional selectivity. Signaling efficacy. Activation mechanism

## Abstract

It has been challenging to determine how a ligand that binds to a receptor activates downstream signaling pathways and to predict the strength of signaling. The challenge is compounded by functional selectivity, in which a single ligand binding to a single receptor can activate multiple signaling pathways at different levels. Spectroscopic studies show that in the largest class of cell surface receptors, 7 transmembrane receptors (7TMRs), activation is associated with ligand-induced shifts in the equilibria of intracellular pocket conformations in the absence of transducer proteins. We hypothesized that signaling through the μ opioid receptor, a prototypical 7TMR, is linearly proportional to the equilibrium probability of observing intracellular pocket conformations in the receptor-ligand complex. Here we show that a machine learning model based on this hypothesis accurately calculates the efficacy of both G protein and β-arrestin-2 signaling. Structural features that the model associates with activation are intracellular pocket expansion, toggle switch rotation, and sodium binding pocket collapse. Distinct pathways are activated by different arrangements of the ligand and sodium binding pockets and the intracellular pocket. While recent work has categorized ligands as active or inactive (or partially active) based on binding affinities to two conformations, our approach accurately computes signaling efficacy along multiple pathways.

## INTRODUCTION

Cell surface receptors transduce extracellular signals into multiple intracellular pathways. Remarkably, in a phenomenon known as functional selectivity^1^ or biased signaling, a single ligand binding to a single receptor can have different effects on distinct signaling pathways. In the largest class of cell surface receptors, 7 transmembrane helix receptors (7TMRs) (traditionally known as G-protein coupled receptors), ligands can differentially activate or inhibit pathways involving heterotrimeric G proteins, 7TMR kinases, and β-arrestins. Signaling efficacy (E_max_ from concentration-response curves) quantifies the extent of pathway activation at a saturating concentration of ligand. For any given pathway, E_max_ can range from near 100% for a full agonist, less for a partial agonist, basal for a neutral antagonist, or even negative for an inverse agonist. Some 7TMR ligands are balanced, with comparable efficacy for both G protein and β-arrestin pathways; others are biased, with much higher efficacy for one class of pathways. While 7TMRs are particularly useful targets for drugs (“druggable”), targeted by approximately one third of drugs in the clinic,^2^ most of these drugs were designed assuming that they would be balanced.^1^ As inappropriate pathway modulation may cause adverse side effects, optimizing functional selectivity is likely to produce safer and more effective drugs targeting 7TMRs^3–6^ and other signaling proteins.

The tragic history of synthetic opioids starkly illustrates the importance of functional selectivity. Fentanyl and its derivatives block pain, exhibiting their analgesic effects by binding to a 7TMR, the μ opioid receptor (MOR). Although the precise pathways are still debated, adverse side effects of tolerance and respiratory depression are also mediated through the MOR.^7–9^ The medicinal chemists who designed the first synthetic opioids reasoned that compounds with high analgesic potency would be safer than morphine.^10^ They touted the high binding affinity of sufentanil to the MOR.^11^ Unfortunately, the hypothesis that potent compounds would be safe was incorrect; due to their dangerous side effects, synthetic opioids have become the leading cause of drug overdose deaths in the United States!^12^

Increased recognition of the importance of functional selectivity has inspired extensive research into its mechanisms.^1^ The focus of the present paper is on ligand-mediated functional selectivity. This type of functional selectivity is independent of mutation or differential splicing of the receptor or differential expression of transducer elements or downstream effectors.^1^ The mechanism of ligand-mediated functional selectivity is generally believed to be stabilization of intracellular pocket conformations that differentially interact with proteins that transduce signals further downstream.^13,14^

Spectroscopic methods show that different classes of ligands have different effects on 7TMR conformational dynamics, even in the absence of transducer coupling.^14^ Wingler et. al. used double electron-electron resonance spectroscopy to show that ligands with different levels of bias can induce at least four sets of conformations of the angiotensin II type 1 receptor.^15^ For the β2 adrenergic receptor, nuclear magnetic resonance (NMR) and single-molecule fluorescence have demonstrated that balanced versus biased ligands have different effects on receptor conformational exchange.^16,17^ Cong et. al. applied NMR to the MOR to show that biased, unbiased, and partial agonists stabilize different conformations of the receptor.^18^ While spectroscopic methods demonstrate the existence of multiple conformations, they have not identified specific three-dimensional structures or determined the extent to which they activate signaling along different pathways.

High-resolution structures provide detailed information about a limited subset of intracellular pocket conformations. X-ray crystallography and cryo-EM structures of 7TMRs are typically solved in complexes that comprise stabilizers, such as antibodies and transducers. These restrict conformational heterogeneity, making structures easy to solve but also obscuring activation mechanisms. MOR structures have been solved as complexes with 17 different ligands^19^ with multiple distinct chemical scaffolds and classes of signaling activity, including partial agonists.^20^ Even though there are a variety of ligands, receptor conformations fall into only two categories: active, for structures complexed to G proteins; and inactive, for structures complexed to antagonists. Presumably the former are capable of G protein signaling while the latter do not activate signaling along any pathway. Even allosteric modulators binding to other 7TMRs do not produce structures with significant changes to high-resolution structures distal to their binding sites.^21^ Two agonist-bound structures of the closely-related δ opioid receptor^22^ may be categorized as intermediate;^19^ they feature outward rotations of helix 5 and 6 and inward rotation of helix 7 indicative of 7TMR activation,^23^ but the tip of helix 6 is less tilted than in the active structures of the μ and κ opioid receptors.^22^ Another 7TMR, the angiotensin II type 1 receptor, has been crystallized in distinct active conformations in complex with balanced versus biased ligands.^24^ These notable exceptions show that is it difficult to capture unique intracellular pocket conformations in high-resolution structures of 7TMRs.

Molecular dynamics simulations (MDS) reveal additional 7TMR conformations. Distinct intracellular pocket conformations have been observed in simulations of MOR complexes with a wide variety of ligands.^18,25–28^ We hypothesized that signaling efficacy is linearly proportional to the equilibrium population of these intracellular receptor conformations. To test this hypothesis, we first performed simulations of MOR complexed with 11 ligands from a variety of chemical series, comprising the most comprehensive set of agonists in a single study to date (previous studies^18,25–28^ included up to 5 complexes). We tested this hypothesis by developing a machine learning model that categorizes configurations from MDS into conformations. Features for the machine learning model are interhelical distances, torsion angles, and hydrogen bonds throughout the intracellular half of the receptor. Fractions of simulation time in each conformation, which are estimates of equilibrium probability, were used as independent variables for multiple linear regression. Model outputs were signaling efficacies along G protein and β-arrestin pathways.

## METHODS

### Molecular Dynamics Simulation

We built three-dimensional models of human μOR bound to 11 ligands based on experimental structures available in the Protein Data Bank (Table S1): FH210, mitragynine pseudoindoxyl, lofentanil, c6guano, c5guano, fentanyl, morphine, TRV130, SR10718, PZM21, and DAMGO. Models were built based on the first chain of the 7TMR that appears in each file, excluding additional 7TMR subunits and G proteins. The apo structure was based on the DAMGO-bound structure 8EFQ with DAMGO removed. Proteins were protonated with pdb2pqr (version 3.6.1)^29^ with a pH of 7.0. Ligands were protonated using RDKit (version 2023.03.1) with a pH of 7.0. Protein-ligand complexes were solvated with 0.15 M NaCl and inserted into a membrane using our group’s custom scripts (https://github.com/swillow/pdb2amber). The scripts build a DPPE lipid bilayer around the protein after alpha carbon alignment to a μOR structure (5C1M) in the Orientations of Proteins in Membranes^30^ database. Complexes were parameterized using AMBER forcefields ff14SB^31^ for the protein, OPC3^32^ for the water, and lipid17 for the membrane. Ligands were parameterized using the GAFF2 force field from AmberTools (version 22.0).^33^

MDS were performed using OpenMM version 8.0.0.^34^ The systems were minimized using the local energy minimizer *simtk.openmm.app.simulation.Simulation.minimizeEnergy* with 500 kJ/mol/nm^2^ restraints on the protein and membrane, 5000 iterations, and a tolerance of 100 kJ/mol. Equilibration was performed in several stages. First, water and membrane were equilibrated with 500 picoseconds of NVT simulation with 300 kJ/mol/nm^2^ restraints on the protein and z-coordinate positions of the membrane. Next, two cycles of 5 nanosecond NPT simulation were performed, with the first cycle using a Monte Carlo Membrane Barostat and the second using a Monte Carlo Barostat. All equilibration simulation was performed with a time second of 2 femtoseconds at 300 K using the Langevin Middle Integrator. Production simulations were performed in triplicate for 500 ns each with a timestep of 3 femtoseconds, saving configurations every 7.5 picoseconds. Production runs were performed at 300 K and 1 bar of pressure with a Monte Carlo Barostat and powered by the Langevin Middle Integrator. Calculations were performed using computing resources provided by the Advanced Cyberinfrastructure Coordination Ecosystem: Services & Support (ACCESS) program.^35^

### Efficacy Calculations

We developed a machine learning model to compute signaling efficacies. Model inputs were configurations from MDS of complexes with each ligand. Outputs were experimental efficacies curated from the literature (Tables S1 and S2). Experimental data from the cyclic adenosine monophosphate (cAMP) assay, which measures the inhibition of downstream cAMP production, were used for G protein signaling efficacies.^36^ For the β-arrestin-2 pathway, we included data from standard assays for measuring β-arrestin-2 recruitment:^36^ NanoBit, BRET, PathHunter, and Tango. Assays performed with G protein receptor kinases were excluded.

The model is particularly simple and interpretable, based on multiple linear regression (MLR). First, configurations from MDS are clustered into *C* conformations. This step yields *f_l,c_*, the fraction of simulations with ligand *l* in conformation *c*. Next, the signaling efficacy is computed based on a weighted sum over all conformations,

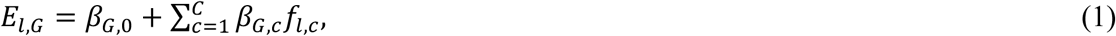

where *E_l,G_* is the signaling efficacy of ligand *l* and *β_G,n_* are regression slopes along G protein pathway. Analogous terms for the β-arrestin-2 pathway, *E_l,β_* and *β_β,n_*, are computed via an analogous expression. We used the MLR implementation in the open source python package scikit-learn (version 1.3.0).^37^

The machine learning model has parameters and hyperparameters. The parameters are regression slopes. Once conformations are defined and fractions computed, these are uniquely defined by the least squares solution for a particular set of input populations and output efficacies. However, the process of defining conformations involves hyperparameters (1) for distances between configurations and (2) for clustering.

#### (1) Distances between configurations

Each sampled configuration was characterized using three vectors of features (Fig. S12). We use Ballesteros-Weinstein nomenclature^38^ in which superscripts describe transmembrane helix (TM) or intracellular loop (ICL), followed by the position relative to the most conserved residue. The vectors are: 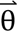, the *N*_θ_ backbone and side chain torsion angles (*ψ*, ϕ, χ_1_, and χ_2_ angles) of all the intracellular protein residues (G84^1.46^-D116^2.50^, S156^3.39^-W194^4.50^, P246^5.50^-F291^6.44^, Y328^7.43^-F349^H8^); 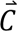, the *N_C_* pairwise distances between alpha carbons from residues in the middle and intracellular end of each helix, with at least one residue in between, and the middle of each intracellular loop (G84^1.46^, V91^1.53^, T99^ICL1^, K102^ICL1^, T105^2.39^, D116^2.50^, S156^3.39^, S164^3.47^, A170^3.53^, L178^ICL2^, P183^4.39^, W194^4.53^, C253^5.57^, R260^5.64^, L267^ICL3^, K271^6.11^, M283^6.23^, F291^6.30^, Y328^7.43^, Y338^7.53^, G343^H8^, and F349^H8^); and 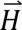, the *N_H_* distances between intracellular hydrogen bond donors and acceptors observed to be within 8.0 Å of the first configuration of FH210-bound complex (PDB ID 7SCG).^39^ For each vector, we use the subscript 𝑘 as an index. The distance between configurations i and j was defined as,

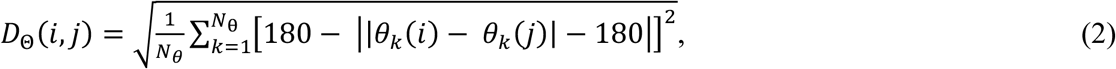

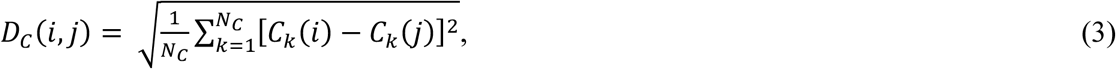

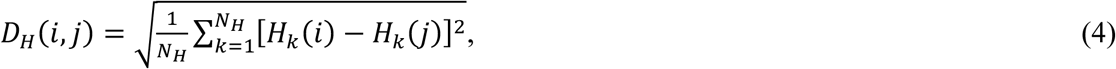

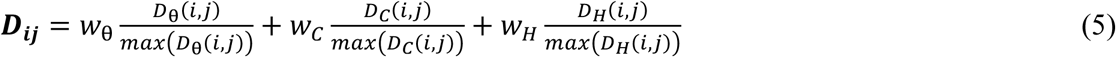

Equation 2 is based on the smallest difference, accounting for periodicity, between torsion angles. Equations 2, 3 and 4 demonstrate how each component was calculated, while equation 5 shows how components are combined to represent a single value for the distance between frames i and j. The sum of weights was constrained to one, such that *w*_θ_+*w_C_* + *w_H_* = 1.

#### (2) Clustering

Intracellular conformations were defined by clustering. First, hierarchical clustering with complete linkage was performed using scipy (version 1.11.3)^40^ based on pairwise distances calculated from equation 5, leading to H hierarchical clusters. Second, the pairwise RMSD between the hierarchical cluster centroids was calculated. The RMSD distance matrix of centroids ***D_rmsd_*** was converted into a similarity matrix ***S_rmsd_*** using a Gaussian function implemented in scikit-learn,^37^

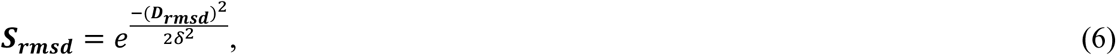

which introduces a bandwidth hyperparameter *δ*. Centroids and corresponding configurations were grouped into conformations using spectral clustering from scikit-learn with the cluster_qr assignment algorithm. This algorithm uses ***S_rmsd_*** and returns C conformations.

### Training and Cross-validation

Leave-one-out procedures were used both for training and cross-validation. In statistics, cross-validation involves separating the data into a training set and a test set. A machine learning model that does not overfit the data produces accurate outputs not only for the training set, but also for the test set. In leave-one-out cross validation, the test set comprises of one sample and the training set is the remainder of the data. The training is repeated multiple times with each sample as the test set.

The machine learning model was trained based on a leave-one-out loss function. Each training set was further divided into a sub-test set comprising one sample and sub-training set containing the remainder of the data. Sub-training was repeated multiple times with each training set sample as the sub-test set. The loss function was the mean square error of the sub-test set, averaged over all the sub-training processes for the given training set.

The loss function was optimized via a grid search. Distance components were weighted according (*w*_θ_, *w*_C_, *w*_H_) ∈ {(*w*, *w*, 1 − 2*w*), (*w*, 1 − 2*w*, *w*), (1 − 2*w*, *w*)}, where *w* ∈ {0.1, 0.2, 0.25, 0.33, 1}. Clustering hyperparameters were varied over a range of integers, with the number of clusters H between 2 and 40, the bandwidth δ between 1 and 3, and the number of conformations C ∈ {2, 3, . . ., *H* − 1}. For each combination of hyperparameters, the leave-one-out loss was computed for both G protein and β-arrestin-2 efficacy. Hyperparameters were selected based on minimizing the sum of the leave-out-out loss of both efficacies.

The machine learning model was validated via leave-one-out cross-validation. Reported signaling efficacies and performance metrics are based on these models which do not include test compounds within training sets. This procedure is consistent with the way efficacy would be predicted for a ligand with a known binding pose but unknown efficacy.

### Structural Analysis

To help us understand the relationship between structural features and receptor activation, we defined an efficacy response function (ERF) based on the defined conformations and MLR weights from the machine learning model. First, we recorded the slopes from each training iteration of the single conformational space that minimizes the loss function. For each structural feature, we computed a kernel density estimate (KDE) of the probability density function within each conformation using numpy.histogram, with the density feature (version 1.26.0).^41^ Each torsional sample Φ was triplicated at Φ + 2π and Φ - 2π to make the density estimate continuous over the period. The KDE was normalized based on evaluating the integral over the range [-π, π] using numpy.histogram, with the density feature (version 1.26.0).^41^ Next, the normalized KDEs were multiplied by corresponding mean MLR slopes and summed together, resulting in efficacy response functions. ERFs were computed for all features that were used to compute distances (see Distances between configurations) and for both G protein and β-arrestin-2 activation. ERFs calculations were extended to residues in the binding pocket, which did not contribute to distances, and weighted with the corresponding intracellular configuration.

An ERF helps identify changes in the probability density that favor an activation process. It is important to note that an ERF is not a probability density function. Because MLR slopes may be negative, an ERF can be negative. Moreover, they are not normalized. Nonetheless, ERFs are helpful for interpreting the machine learning model. Shifting a probability density towards a region with high ERF values favors activation. Conversely, shifting a probability density towards a region with low ERF values favors inactivation. If the ERF is near zero over the entire range, the feature is unrelated to activation.

As we computed many ERFs, we also defined several metrics to help us prioritize, in an unbiased way, which structural features to focus our attention on. We defined the general activation function *a*(*x*) as,

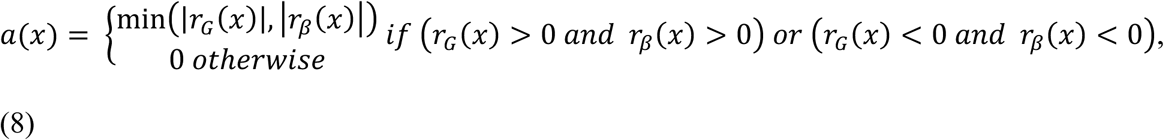

where *r_G_*(*x*) and *r_β_* are ERFs for the G protein and β-arrestin-2 pathways, respectively. This function is nonzero in regions where ERFs of both pathways have the same sign. Conversely, the selective activation function is,

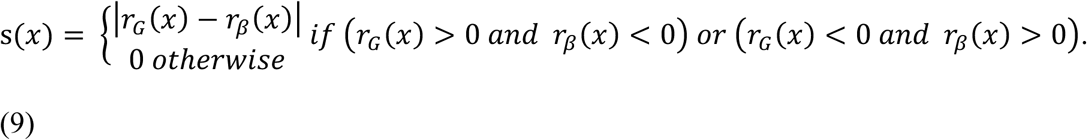

This function is nonzero in regions where ERFs of both pathways have opposite signs. For each function, we also defined corresponding scores as a numerical integral over the domain: the sum of the function was computed 1,000 evenly spaced points over the domain and multiplied by the spacing.

## RESULTS

### Leave-one-out training leads to consistent values of all machine learning hyperparameters

Leave-one-out training led to consistent values of all machine learning hyperparameters. For all 11 models trained with each ligand as the test set, the best performance was observed with the distance weights of *w*_θ_ = 0.25, *w*_C_ = 0.25, and *w*_H_ = 0.5, the number of hierarchical clusters H = 40, and bandwidth δ = 2, and the number of conformations C = 14. The consistency of these parameters indicates that the training procedure is robust, insensitive to the inclusion or exclusion of any single ligand. Thus, these hyperparameters were used for all models reported in the paper.

### The machine learning model accurately predicts signaling efficacy along two pathways

Model predictions of E_max_ are accurate and precise (Fig. 1). Compared to median experimental values, the efficacy of G protein and β-arrestin-2 (βarr2) pathways is predicted with a mean absolute error of 8.1% and 18.4% and a root mean squared error of 10.8% and 21.2%, respectively. The coefficients of determination (*R*^2^) are 0.67 and 0.69, respectively. The standard deviation of efficacy predictions is small, a demonstration that the training procedure is robust.

**Figure 1.**
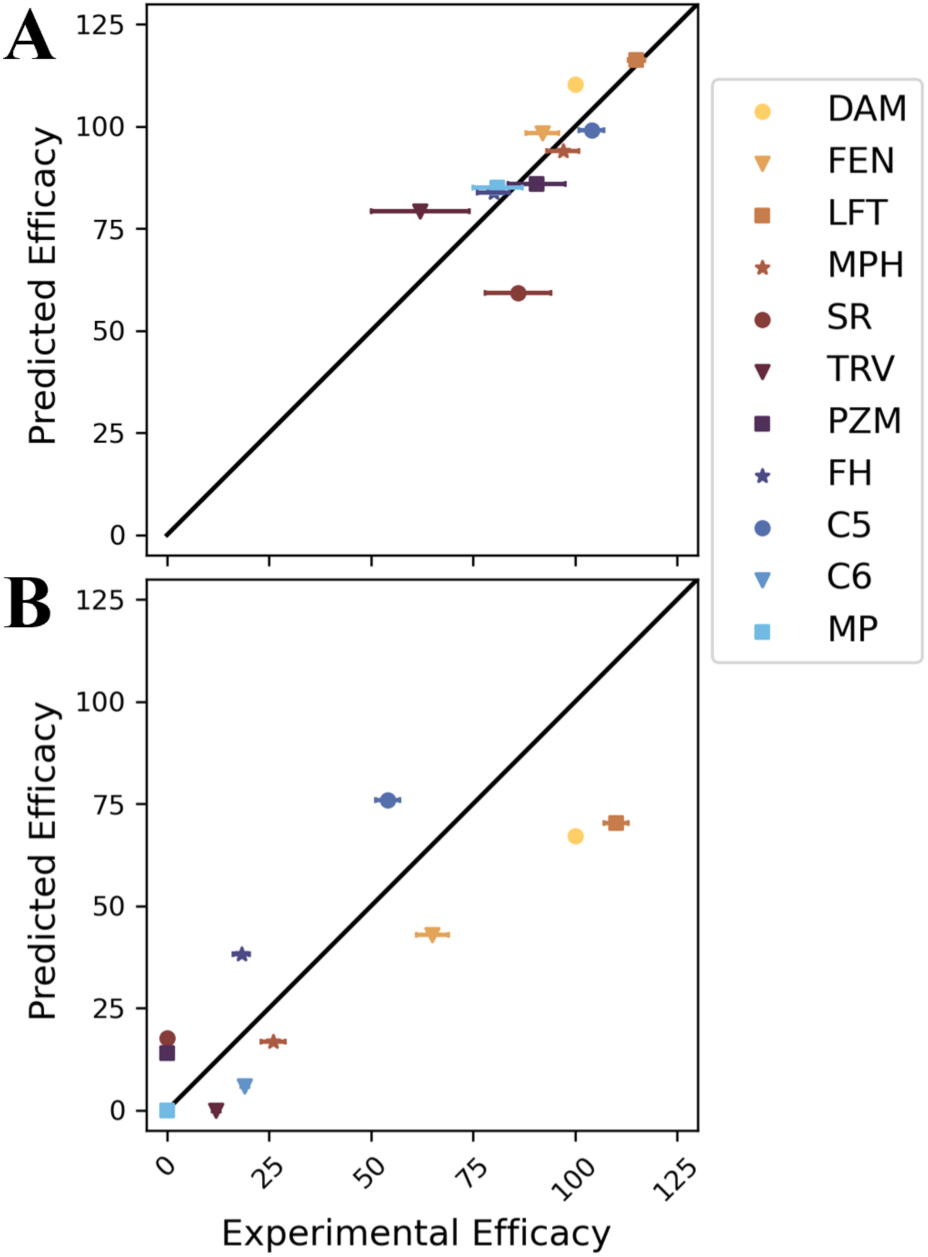
Accuracy of signaling efficacy predictions of ligands for (A) G protein and (B) βarr2 pathways. Efficacies are percentages relative to DAMGO. On the x axis, each experimental efficacy is the median of reported values listed in Table S1. Error bars are standard deviations of these values. On the y axis, each predicted efficacy is based on a model trained using all other ligands. The error bar is the standard deviation across 11 cross validation models. Abbreviations: DAM: DAMGO, FEN: fentanyl, LFT: lofentanil, MPH: morphine, TRV: TRV130, SR: SR17018, PZM: PZM21, FH: FH210, C5: c5guano, C6: c6guano, MP: mitragynine pseudoindoxyl.

The performance of models trained with different lengths of simulation is also robust (Fig. S1). While the mean square error fluctuates as the length of simulation is varied from 100 to 500 ns (in triplicate) per protein-ligand complex, it remains in the general range of 10% to 20%. Further discussion is based on a model with 500 ns.

The model is based on 14 conformations with a wide range of activity (Fig. 2). We indexed these conformations in decreasing order by average regression slope along G protein and βarr2 pathways; conformation 1 has the largest average slope and conformation 14 the smallest. For this reason, we used these two conformations to represent active and inactive conformations, respectively, in Fig. 3. However, the average oversimplifies the activity of these conformations. Conformations 4 and 9 promote G protein signaling and repress βarr2 signaling. Conformations 2, 3, 5, and 7 promote βarr2 signaling at different levels but have minimal effect on G protein signaling. For this reason, we used conformation 4 to visualize G protein signaling and conformation 3 to visualize βarr2 signaling. Conformations 1 and 6 recruit both G proteins and βarr2. The remaining conformations repress signaling.

**Figure 2.**
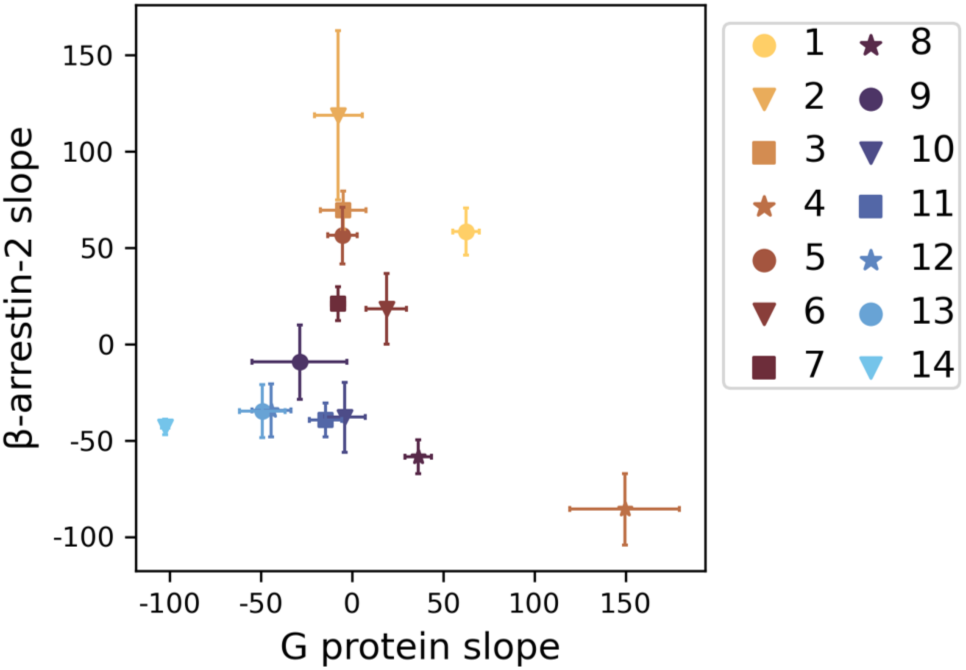
Multiple linear regression slopes of each conformation. The mean (markers) and standard deviation (error bars) are of slopes across all 11 cross validation models.

**Figure 3.**
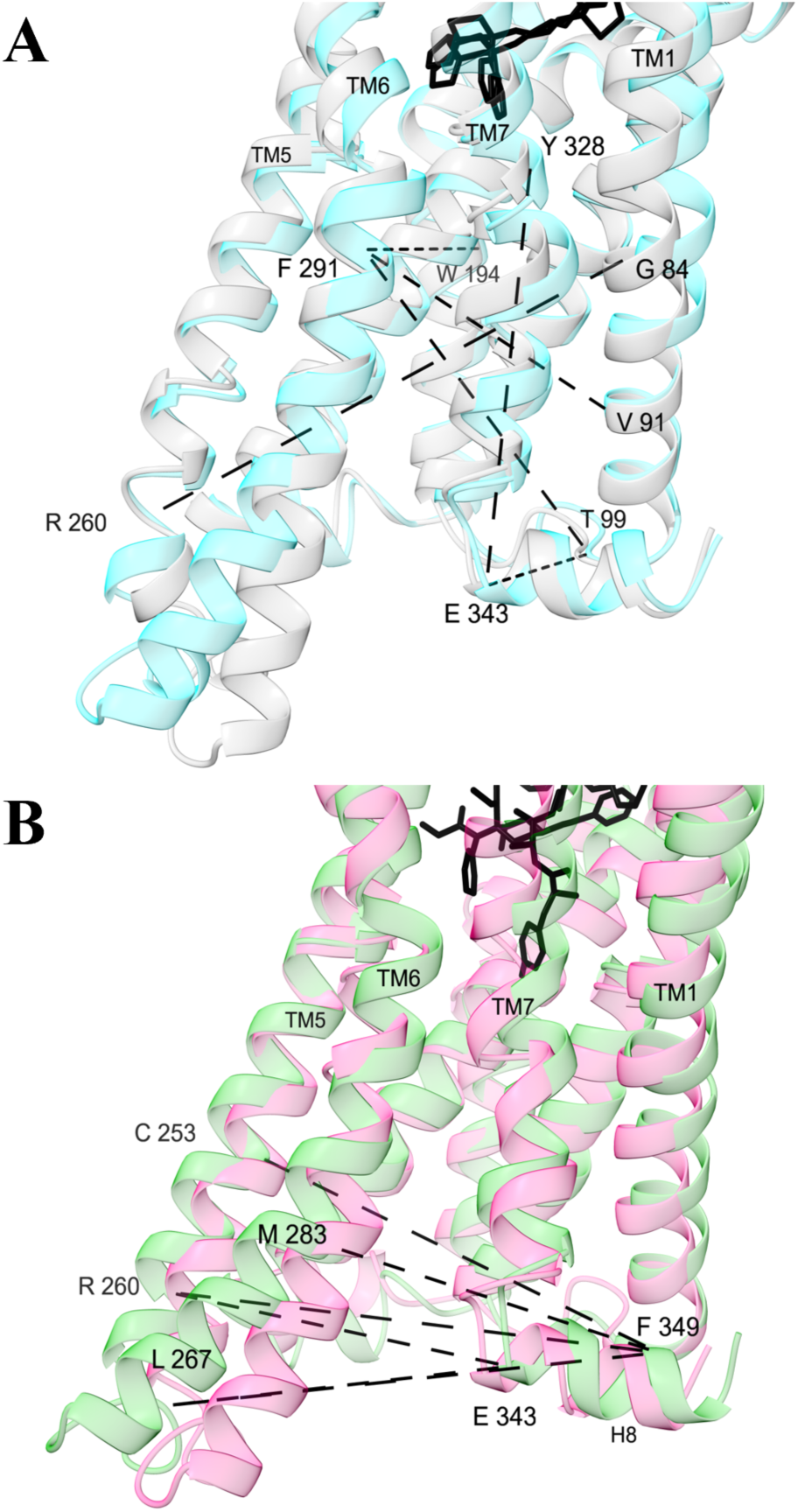
Alpha carbon distances with the largest (A) general and (B) selective activation scores. Structures are medoids of conformations from the machine learning model, colored by different types of activity: high (cyan, conformation 1), low (grey, conformation 14), G protein (green, conformation 4), and βarr2 (pink, conformation 2). They are shown from the membrane perspective facing the intracellular pocket. Dashed lines are between atoms are shown for the top 2% of activation function scores. For a full table of general and selective activation scores for alpha carbon distances, see Fig. S6.

Simulations with different ligands access different ratios of these shared conformations (Fig. S2). Some simulations, such as the apo system (conformation 11) and the complex with DAMGO (conformation 4), are dominated by a single conformation. Others, such as simulations of complexes with PZM21, c5guano, and c6guano, are spread across two or more conformations. Complexes with comparable ligands primarily access distinct conformations, explaining differences in efficacy. Complexes with fentanyl and lofentanil both access conformation 6, but it is not the most populated conformation of either. Similarly, conformation 8 is shared between complexes with both c5guano and c6guano, but both complexes are dominated by other conformations.

Most conformations are accessed in simulations with multiple ligands (Fig. S3). However, there are three conformations that are unique to complexes with a specific ligand: 2 (lofentanil), 12 (fentanyl), 13 (FH210).

### Signaling is associated with specific structural features

We defined several functions to help us understand the relationship between structural features and receptor activation (see Materials and Methods). The efficacy response function (ERF) is a sum of estimated probability density functions of a structural feature weighted by linear regression slopes. The general and selective activation scores quantify whether the ERF is significant in both or only one of G protein and βarr2 recruitment.

Activation is associated with rearrangement of helices in the intracellular pocket (Fig. 3A). In structures that favor activation, the intracellular end of TM5 is bent closer to TM1, such that the G84^1.46^-R260^5.64^ distance is reduced by 2 to 4 Å (Fig. S4A). TM6 is pushed outwards away from TM1, increasing the V91^1.53^-F291^6.30^ and T99^ICL1^-F291^6.30^ distances (Fig. S4BC). A kink above a proline in TM7 becomes stronger such that the Y328^7.43^-E343^H^^8^ distance is reduced (Fig. S4F). Compared to βarr2 activation, G protein activation is favored when TM5 and TM6 are bent further outward relative to helix 8 (Fig. 3B, Fig. S5). Activation is also associated with side chain dihedral angles of several amino acid residues in the orthosteric binding pocket (Fig. 4, Fig. S7). In W295^6.48^, activation relates to a shift in the χ_2_ angle from around - 120° to around 120° (Fig. S7J), rotating the indole ring towards the intracellular pocket. Across the binding pocket, D149^3.32^,

**Figure 4.**
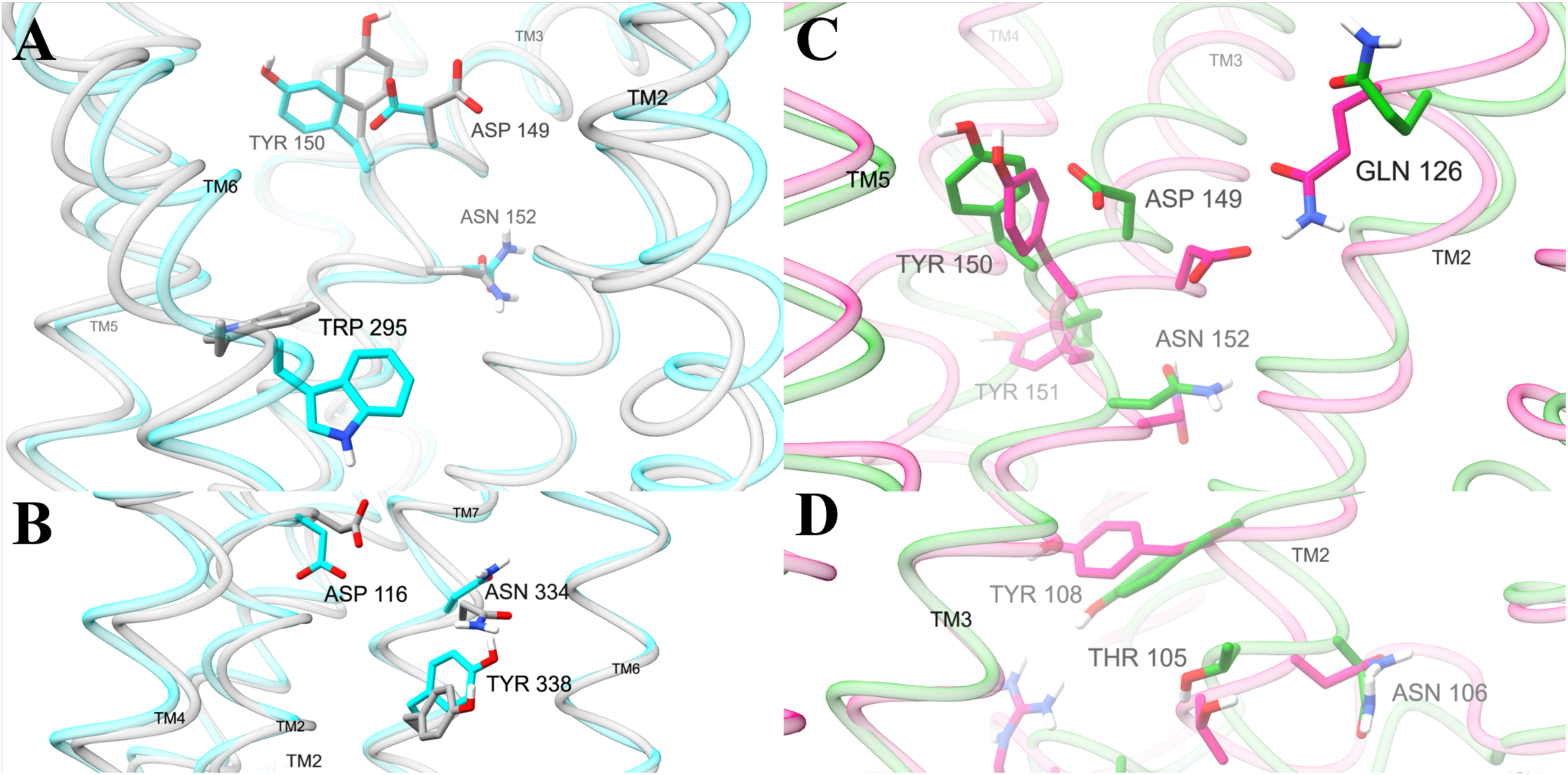
Amino acid positions containing dihedral angles with the highest general and selective activation scores. Structures are medoids of conformations from the machine learning model, colored by different types of activity: high (cyan, conformation 1), low (grey, conformation 14), G protein (green, conformation 4), and βarr2 (pink, conformation 2). Displayed amino acids contain dihedral angles in the top two percent of general (**A**, **B**) activation scores or (**C**, **D**) selective activation scores. Side chain rotations were adjusted to match the degree with the highest value of the (A, B) general or (C, D) selective activation function. Views of the binding pocket (A, C) are from the perspective behind the binding pocket region of TM7. TM1 and TM7 (A) and TM6 and TM7 (C) were removed to improve the display of binding pocket amino acids. Views of the intracellular region are from the perspective behind TM3 (B) and TM7 (D). TM3 (B), TM6 and TM7 (D) are removed to clearly display the intracellular amino acids. A comprehensive table of general activation and selective activation function scores for analyzed residues are shown in Fig. S9 and S10, respectively.

Y150^3.33^, and N152^3.35^ have distinct active and inactive conformations. In active conformations, the carboxylate of D149^3.32^ and the phenol of Y150^3.33^ extend across the binding pocket towards TM5 and TM6. In contrast, D149^3.32^ tucks towards TM2 and Y150^3.33^ extends back towards TM4 in inactive conformations. The orientation of N152^3.35^ coincides with D149^3.32^ and Y150^3.33^, pointing the side chain amide further away from the center of the receptor in active conformations (Fig. 4A). D149^3.32^ and Y150^3.33^ also have high selective activation scores; in G protein selective conformations, the side chains of these residues are further oriented towards a sub pocket of the binding site between TM3, TM4, and TM5 (Fig. 4C).

Several residues in the interior of the intracellular pocket also have distinct orientations in active and inactive conformations (Fig. 4, Fig. S7). D116^2.50^ extends down towards intracellular space in active conformations opposed to upwards towards interhelical space in inactive conformations. Along the interior of intracellular TM7, N334^7.49^ and Y338^7.53^ are rotated upwards in active compared to inactive conformations. At the intracellular interface, polar residues M283^6.23^, D342^H8^, and N344^H8^ assume different side chain configurations in active and inactive conformations.

While intracellular pocket side chain dihedrals with large general activation scores are primarily in TM6, TM7, and H8, those with large selective activation scores are across the pocket in ICL2 and TM2 (Fig. 5, Fig. S8). The alcohol group of T105^2.39^ and the amide group of N106^2.40^ extend towards ICL2 in G protein selective conformations but towards ICL1 in βarr2 selective conformations. Y108^2.42^ orients across the intrahelical space towards TM3 in βarr2 selective conformations but downwards towards intracellular space in G protein selective conformations. On ILC2, R181^ICL2^ reaches into intracellular space in the G protein selective conformations but is retracted into the receptor in βarr2 selective conformations, accompanied by different configurations of nearby L178^ICL2^.

**Figure 5.**
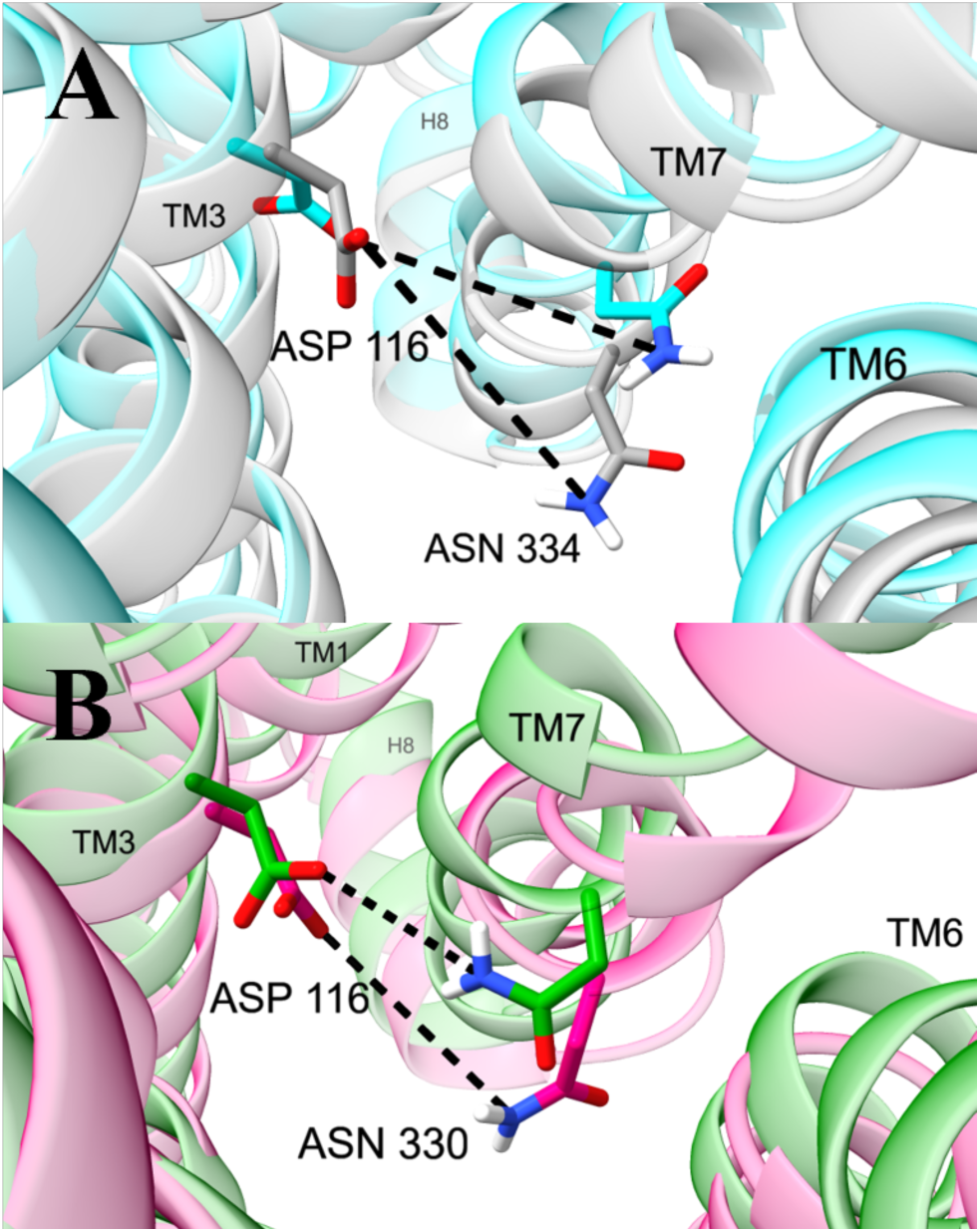
Distances in the sodium binding pocket with the highest general and selective activation scores. Structures are medoids of conformations from the machine learning model. **(A)** Distances between OD1 of D116^2.50^ and ND2 of N334^7.49^ in structures with high (cyan, conformation 1) and low (grey, conformation 14) activity. **(B)** Distances between OD2 of D116^2.50^ and ND2 of N330^7.49^ in conformations with high G protein (green, conformation 4) and βarr2 (pink, conformation 2) activity. Efficacy response functions for analyzed polar residue distances are shown in Fig. S11.

Polar networks in the sodium binding pocket are important both for general and selective activation. The network in this region includes distances between polar atoms of residues D116^2.50^, N330^7.45^, and N334^7.49^ (Fig. 5). Two distances of ∼3 and ∼4 Å between the first carboxylate atom (OD1) of D116^2.50^ and the nitrogen atom (ND2) of the amide side chain in N334^7.49^ are correlated with general activation. The distance between the second carboxylate atom (OD2) of D116^2.50^ and the nitrogen atom (ND2) of the amide side chain of N330^7.45^ differs in G protein (peaked around ∼5.5 Å) opposed to βarr2 signaling (peaked around ∼6.5 Å) (Fig. S11). It is also noteworthy that the neighboring residues W295^6.48^ and N152^3.35^ have dihedral angles with large general activation scores (Fig. S7).

## Discussion

### µOR conformations produce a range of signaling efficacy

The accuracy of our model (Fig. 1) supports the hypothesis that signaling efficacy is proportional to the equilibrium probability of observing intracellular pocket conformations in the receptor-ligand complex in the absence of transducer molecules. Previous observations are also consistent with this hypothesis. For example, Miao and McCammon performed Gaussian accelerated MDS of the muscarinic M_2_ receptor in complex with either an inverse agonist, partial agonist, or antagonist.^42^ They observed that the inverse agonist stabilized an inactive conformation. In contrast, the partial and full agonist stabilized two different intermediate conformations at different levels. Based on our current understanding, the distinct equilibrium probabilities of these intermediate conformations could explain the distinct signaling efficacies of the agonists. In analyzing MDS of the angiotensin II type 1 receptor, Suomivuori et. al. showed that the agonist-bound receptor can transition between two active conformations that are common across complexes with different ligands.^43^ One of the active conformations accommodates both G proteins and β arrestins, but the other favors β arrestin binding. They found that the fraction of simulation time spent in each conformation was related to whether the bound ligand was balanced or biased towards one class of pathways. They also observed three other conformations and speculated about their signaling profiles. We have built upon the concept of multiple conformations with distinct signaling profiles to develop a quantitative model that predicts signaling efficacy.

To our knowledge, we have presented the first computational model that connects conformational equilibria to functional selectivity. There have been two recent attempts to use MDS to categorize functional response without regard to the pathway. Panel et. al. computed relative binding free energies between 23 ligands and active and inactive conformations of the β2 adrenoceptor.^44^ The calculated shift in binding affinity was used to successfully classify the compounds as agonists, partial agonists, or antagonists. In another study, a team from the computational chemistry software company Schrödinger performed absolute binding free energy calculations of ligands to active and inactive conformations of 7TMR and nuclear hormone receptor targets.^45^ Based on higher predicted binding affinity to the active conformation, 168 of 180 ligands (93%) could be correctly classified as agonists opposed to antagonists. We have not only categorized functional response as on or off (or partially on) but provided a quantitative prediction of efficacy and evaluated the ability of different conformations to activate signaling along multiple pathways.

In the context of the previous state of the field, the accuracy of our model is an important step forward. However, there is significant room for improvement. Two of the clearest future directions to increase the accuracy of our model are enhanced conformational sampling and more consistent training data.

Enhanced conformational sampling is likely necessary to obtain more precise results. In simulations of *agonist-bound* β_2_ adrenergic receptor, Dror et. al. observed that transitions to inactive intracellular pocket conformations can occur on the timescale of several microseconds.^46^ The fact that we did not observe such dramatic conformational changes may be due to our relatively limited sampling or system-dependent differences in the free energy profile, e.g. a higher barrier between active and inactive conformations or a higher free energy of the inactive conformation. Given the longer time scale of the previously observed conformational transitions, it is unlikely that our much shorter simulations (500 ns) have truly converged. Nonetheless, the consistency of machine learning performance across different simulation lengths (Figure S1) suggests that our simulation protocol performs adequate sampling of intracellular pocket conformations capable of capturing transducer molecules. In future studies, enhanced sampling methods such as replica exchange molecular dynamics, which has been successfully applied to 7TMR systems,^43^ could lead to more precise estimates of intracellular pocket populations without a prohibitive increase in computational cost, leading to more accurate efficacy calculations.

Our model could also be improved with a larger and more consistent training set. While additional ligands could lead induce additional conformations, they could also lead to more precise regression slopes. Due to limited availability, not all efficacy data were based on the same assays (Table S2). A model trained on consistent data would likely lead to more accurate efficacy predictions.

Other possible improvements to our model include extension to compute other properties. Other properties that may be proportional to equilibrium populations of intracellular pocket conformations include efficacies of Gα subtypes and parameters of the Black-Leff^47^ operational model: the transducer ratio and dissociation constant of the agonist-receptor-transducer complex.

Our results affirm that many intracellular pocket conformations defy simple classification.^13,43^ As also observed by Miao and McCammon^42^ for the muscarinic M_2_ receptor, ligand-bound receptor conformations do not exactly match those of ternary complexes and cannot be simply classified as active or inactive. Neither are they completely biased towards G protein or β arrestin signaling. Instead, conformations have a broad range of signaling efficacy across multiple pathways (Fig. 2).

While our model identifies conformations and structural features associated with activation, it does not explain how intracellular pocket conformations lead to distinct signaling efficacies. As noted in the introduction, high-resolution structures of 7TMRs in ternary complexes have been observed to be remarkably consistent, regardless of whether ligands are partial or full agonists or whether they are orthosteric or allosteric^21^ and whether transducers are G proteins or βarrs. For G protein pathways, efficacy (which can depend on the specific Gα subtype)^48^ is likely determined by how ligands affect the dynamics of the complexed heterotrimeric G protein to promote the release of GDP and uptake of GTP. For βarr pathways where efficacy is measured by enzyme complementation assays, different ligands can have different effects on catalytic rates. In either case, the conformational equilibria of the receptor-ligand complex without transducer appears to be correlated with how ligand binding perturbs the conformational equilibria of the transducer in the ternary complex.

It has been suggested the signaling efficacy of 7TMR agonists is related the kinetic context.^49^ Strong correlations have been observed between residence time and the efficacy of sets of muscarinic M_3_^50^ and adenosine A_2A_^51^ receptor agonists. However, comparable correlations have been not been observed in studies with adenosine A_1_,^52^ dopamine D_2_,^53^ and cannabinoid CB_2_^54^ receptor ligands. Moreover, MOR ligands have been observed to have high efficacy for some pathways and low efficacy for others.^28^ For example, morphine has high G protein but low βarr efficacy, contrasting with C11 guano which has high βarr2 and G_z_ but low G_i/o_ efficacy. As the same ligand has the same residence time at the receptor, residence time cannot explain these opposite behaviors. Given that efficacy may be time-dependent,^53^ our proposed relationship between intracellular pocket conformation populations and efficacy should be evaluated at different time points, as well as in additional systems.

### Mechanisms of signaling activation are consistent with previous studies

Activation-induced changes in the overall architecture of intracellular pocket conformations predicted by our model are corroborated by previous studies. As we have observed (Fig. 3), high-resolution structures and spectroscopy have shown that activation is favored by expansion of the intracellular recruitment site through rearrangement and bending of the transmembrane helices.^13,55,56^ Moreover, our finding that larger distances between TM5/6 and TM7/H8 are associated with G protein bias is consistent with previous simulations.^26,57^

Orthosteric binding site features that our model associates with functional selectivity are consistent with the pharmacophore model from Kelly et al.^58^ Molecular modeling methods have been used to design several G protein biased ligands.^26,28,39,59^ From an analysis of the interactions between functionally selective agonists and the MOR, Kelly et al. proposed a pharmacophore model in which strong interactions with D149^3.32^ and Y328^7.43^ increase the recruitment of β-arrestin, while interactions with Y150^3.33^ (instead of Y328^7.43^) and a weaker interaction with D149^3.32^ decrease β-arrestin recruitment.^58^ Based on this insight, Zhuang et al. created G protein selective fentanyl derivatives.^26^ Our observation that D149^3.32^ and Y150^3.33^ extend into the binding pocket in active conformations and retract in inactive conformations suggests that ligand interactions with D149^3.32^ and Y150^3.33^ can have an important role in activation (Fig. 4A). Moreover, our observation that D149^3.32^ and Y150^3.33^ point towards the center of the pocket in βarr2 selective conformations suggest that they form stronger interactions with ligands, as suggested by Kelly et al.^58^ In another 7TMR, the apelin receptor, mutagenesis studies demonstrate that I109^3.32^ and F110^3.33^ are part of hot spots that determine signaling bias.^60^ While their ligands have different interactions, β-arrestin bias is associated with weaker interactions between TM3 and TM5 (Fig. S4H-J of Wang et al ^60^), consistent with our observation that Y150^3.33^ in βarr2 selective conformations points away from TM5 (Fig. 4C).

Our observations about the sodium binding site and its allostery are consistent with previous studies. The sodium binding site,^61^ which houses the cation in inactive conformations, comprises D116^2.50^, S156^3.39^, N330^7.45^, S331^7.46^, and N334^7.49^. The site allosterically communicates with the orthosteric site and the intracellular pocket.^13,55,61,62,60^ Ligand-interactions with D149^3.32^ and Y150^3.33^ in the orthosteric site affect the position of the neighboring residue N152^3.35^. In turn, W295^6.48^ and N152^3.35^ form a network with the sodium binding pocket.^63^ These previous studies are consistent with our findings that W295^6.48^ and N152^3.35^ are associated with activation (Fig. S7) and that several polar residues that occupy the sodium binding pocket (D116^2.50^, N330^7.44^, and N334^7.48^) adopt different orientations for generally active, inactive, and selectively active conformations (Fig. 4 and 5). For example, in the apelin receptor, structures bound to G protein biased ligands include a hydrogen bond between D75^2.50^ and N305^7.49^.^60^ Likewise, we observe a hydrogen bound between the homologous residues D116^2.50^ and N330^7.49^ in the MOR (Fig. 5B).

Our analysis predicts behavior of conserved motifs in the MOR that are consistent with previous observations. Three motifs conserved across many 7TMRs – the transmission switch (CWxP from C^6.47^ to P^6.50^, sometimes called the rotamer toggle switch), tyrosine toggle switch (NPxxY from N^7.49^ to Y^7.53^), and ionic lock (DRY including D^3.49^, R^3.50^, and Y^5.58^) – are widely considered microswitches involved in activation.^64^ W295^6.48^ of the transmission switch and N334^7.49^ and Y338^7.53^ of the toggle switch have distinct conformations and large general activation scores (Fig. 5). The rotation of W295^6.48^ has also been associated with activation in other simulations.^18,65^ In contrast, DRY motif residues have significantly smaller activation scores (Fig. S9 and S10). While the arginine in the motif typically coordinates to D/E6.30 in other 7TMRs, MOR has a threonine at this position. Thus, the DRY motif cannot form an ionic contact that stabilizes inactive conformations of other 7TMRs.

Our analysis predicts that some residues at the intracellular end of TM2 (T105^2.39^, N106^2.40^, and Y108^2.42^) and ICL2 (L178^ICL2^ and R181^ICL2^) have distinct configurations in functionally selective conformations. Based on simulations of ternary complexes, Mafi et. al. also proposed that R181^ICL2^ and T105^2.39^ play a key role in recruitment ^57^. The relevance of R181^ICL2^ to G protein signaling and βarr2-mediated internalization has been validated by mutagenesis.^66^ Our analysis suggests that the extension of R181^ICL2^ into cytoplasmic space may increase the affinity for G proteins while higher placement of the polar side chain may improve affinity for βarr2.

### Efficacy prediction may be helpful for drug design

Drug developers are increasingly pursuing functional selectivity of 7TMRs to maximize therapeutic potential and minimize adverse effects. These efforts have been hindered by limited understanding of the structural mechanisms of functional selectivity and the inability to quantitatively predict signaling along multiple pathways. Our approach addresses both challenges and could help design novel functionally selective agonists for the MOR. It may be a platform technology that can be extended to other pathways, such as G protein subtypes, and other signaling proteins.

#### Conclusions

Signaling efficacy of the MOR is linearly proportional to the equilibrium population of intracellular pocket conformations. Equilibrium populations may be accurately estimated by molecular dynamics simulations. Suitable definitions of these intracellular pocket conformations may be determined by training a machine learning model. Intracellular pocket conformations have a broad range of signaling efficacy along different pathways. Efficacy response functions and activation scores are effective metrics for identifying structural features associated with general and selective activation. These analyses lead to predicted structural mechanisms for general and selective activation that are supported by previous computational and experimental studies.

## ASSOCIATED CONTENT

### Data and Software Availability Statement

Signaling efficacy data used for training machine learning models are included in Tables S1 and S2. Initial configurations for molecular dynamics simulations are available on GitHub (https://github.com/CCBatIIT/MOR-Paper-Input-Structures/). Machine learning software implementing the described methods is available through a paid license or collaboration with the authors, who are currently affiliated with the Illinois Institute of Technology and Biagon Inc.

### Supporting Information

Dependence of mean absolute error and the number of conformations on the simulation length, percentages of simulations of complexes with different ligands in each conformation, percentages of conformations accessed in simulations with each complex, efficacy response functions and scores for selected alpha carbon distances, dihedral angles, representations of features used to discretize configurations into conformations, and structures and efficacy data used for MDS and model training.

## Author Contributions

Conceptualization: DDLM, DAC, JDB, SN, BG

Methodology: DDLM, DAC, JDB

Investigation: DDLM, DAC

Visualization: DDLM, DAC

Funding acquisition: DDLM

Project administration: DDLM

Supervision: DDLM

Writing – original draft: DAC, DDLM

Writing – review & editing: DDLM, DAC, JDB

## Notes

DAC and DDLM are co-inventors of the methodology described in this work and the Illinois Institute of Technology has filed a provisional patent on their behalf. DAC, JDB, and DDLM have started a company that is commercializing the technology, Biagon Inc.

## ACKNOWLEDGMENT

We thank Bob Eisenberg for comments on the manuscript.

## Supporting Information

Figures S1 to S12

Tables S1 to S2

SI References

**Figure S1.**
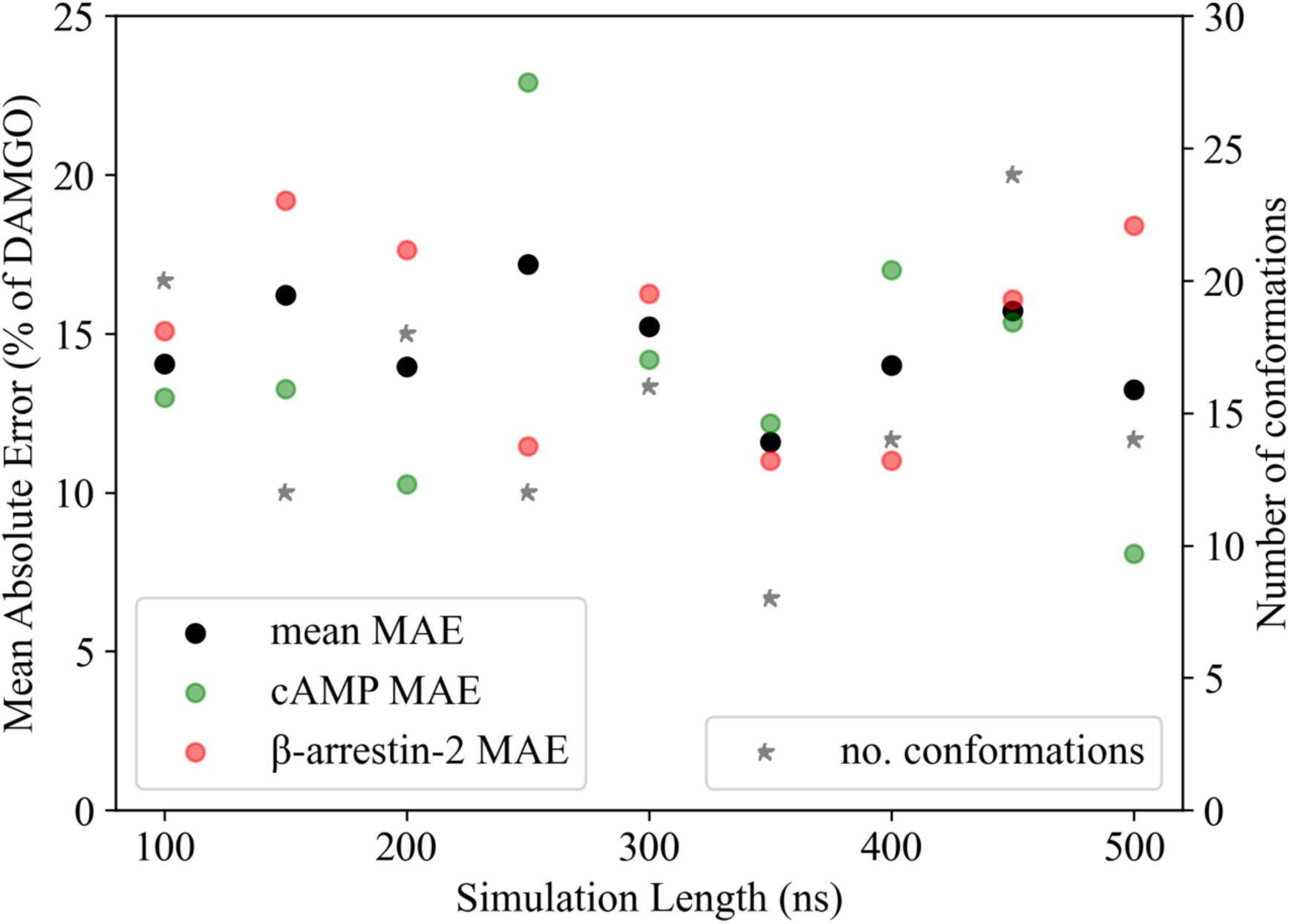
Model parameters and performance as a function of simulation length.

**Figure S2.**
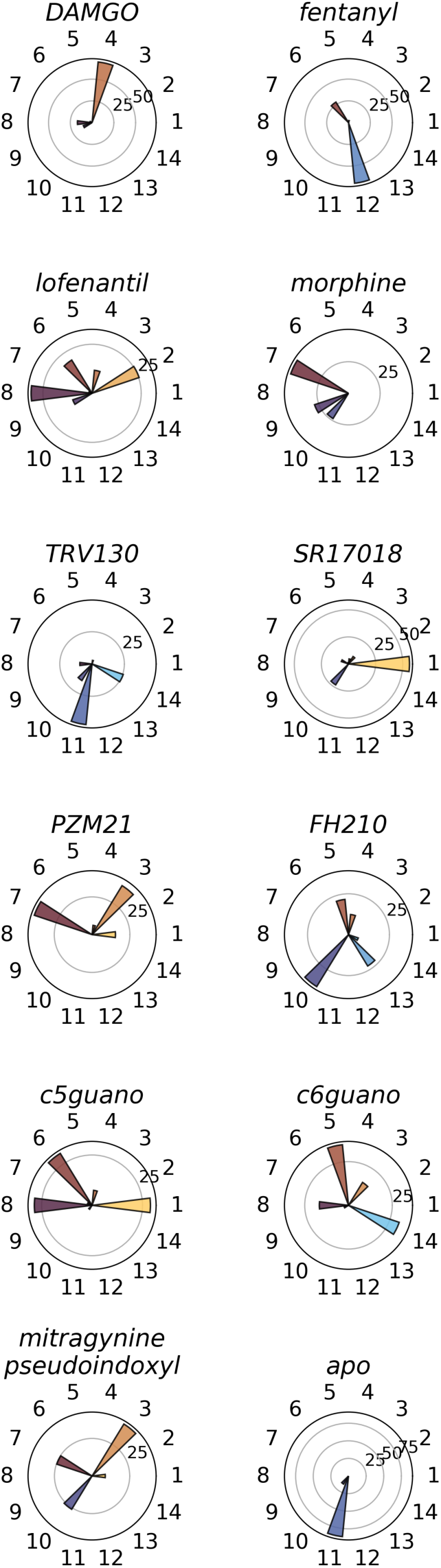
Percentages of simulations of complexes with different ligands in each conformation.

**Figure S3.**
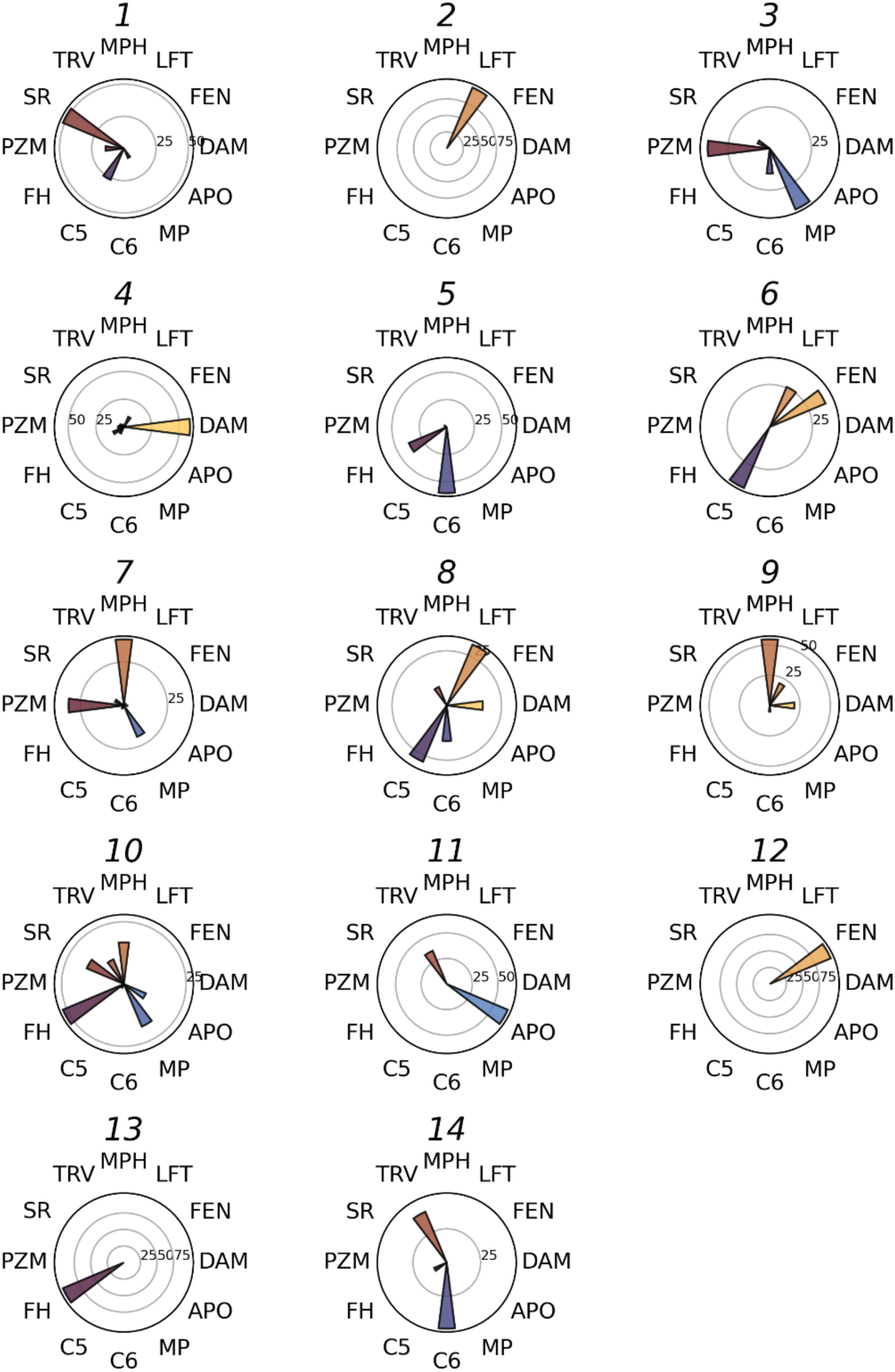
Percentages of conformations accessed in simulations with each complex. Abbreviations: DAM: DAMGO, FEN: fentanyl, LFT: lofentanil, MPH: morphine, TRV: TRV130, SR: SR17018, PZM: PZM21, FH: FH210, C5: c5guano, C6: c6guano, MP: mitragynine pseudoindoxyl, APO: apo.

**Figure S4.**
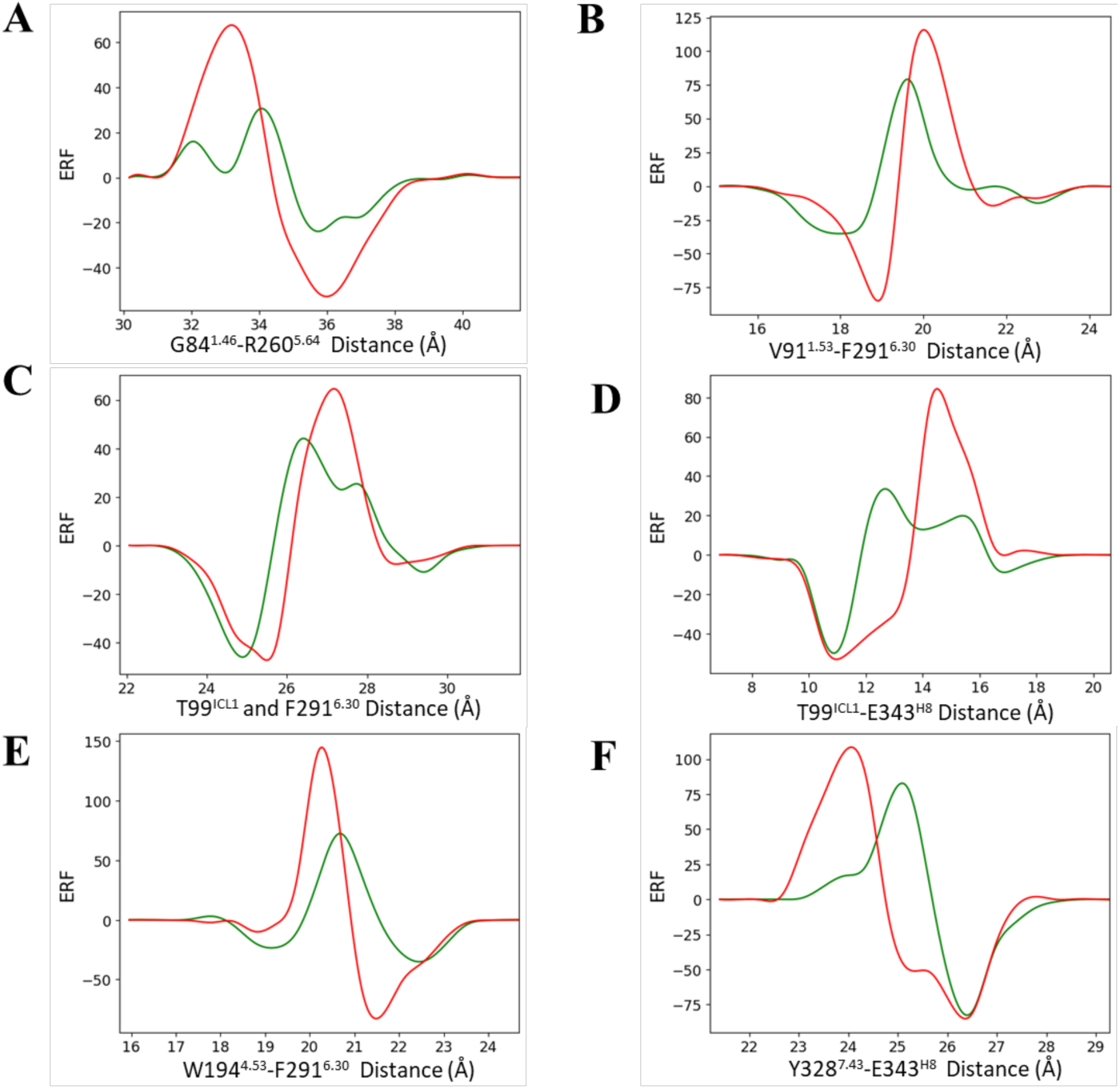
Efficacy response functions for alpha carbon distances with the largest general activation scores: (A) G84^1.46^-R260^5.64^, (B) V91^1.53^-F291^6.30^, (C) T99^ICL1^-F291^6.30, (D) T99ICL1-E343H8, (E) W1944.53-F2916.30, and (F) Y3287.43-E343H8. The green and red curves correspond to the G protein and β-arrestin-2 efficacy response functions (ERF), respectively.^

**Figure S5.**
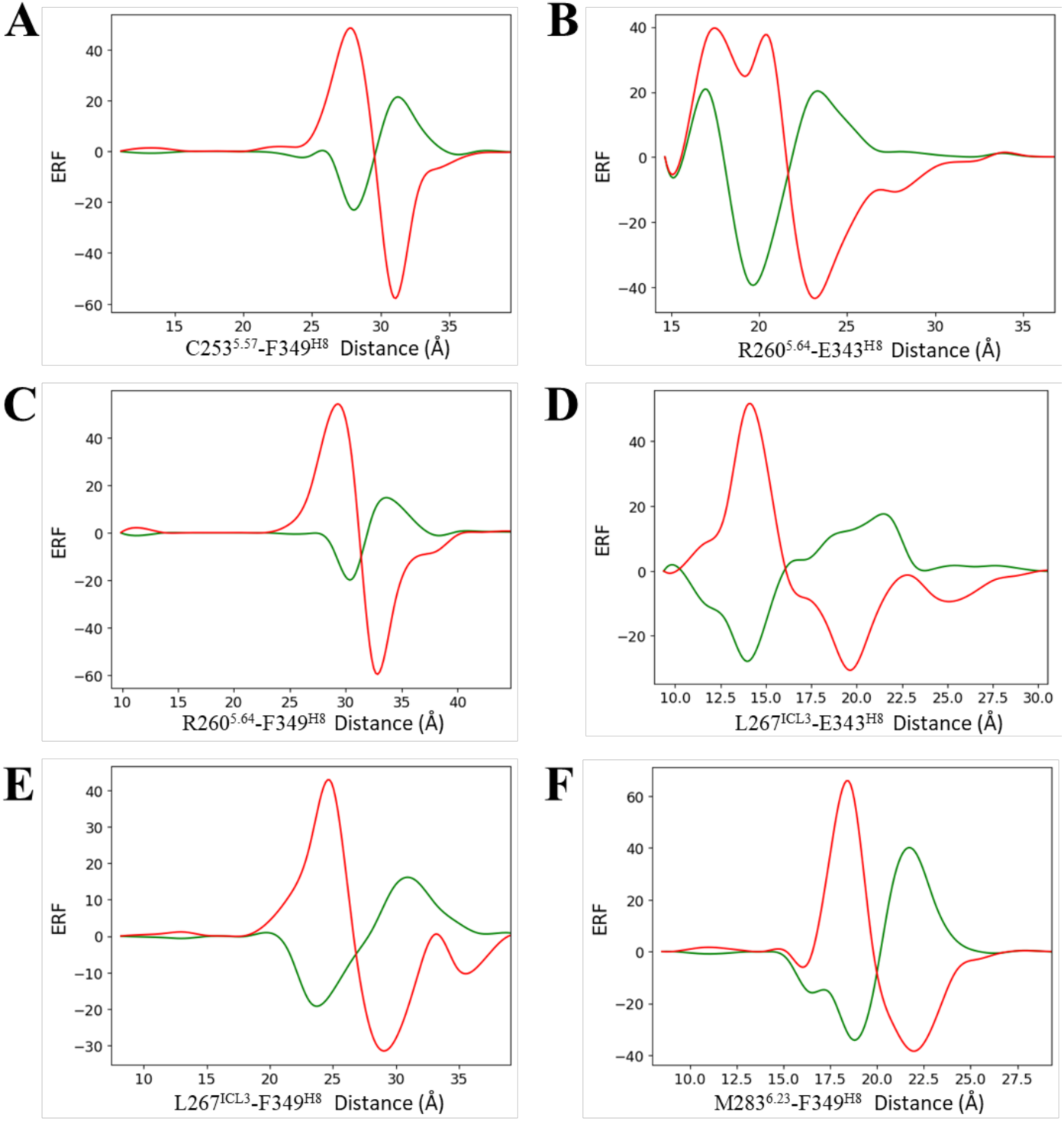
Efficacy response functions for alpha carbon distances with the largest selective activation scores: (A) C253^5.57^-F349^H8^, (B) R260^5.64^-E343^H8^, (C) R260^5.64^-F349^H8^, (D) L267^ICL3^-E343^H8^, (E) L267^ICL3^-F349^H8^, and (F) M283^6.23^-F349^H8^. The green and red curves correspond to the G protein and β-arrestin-2 efficacy response functions (ERF), respectively.

**Figure S6.**
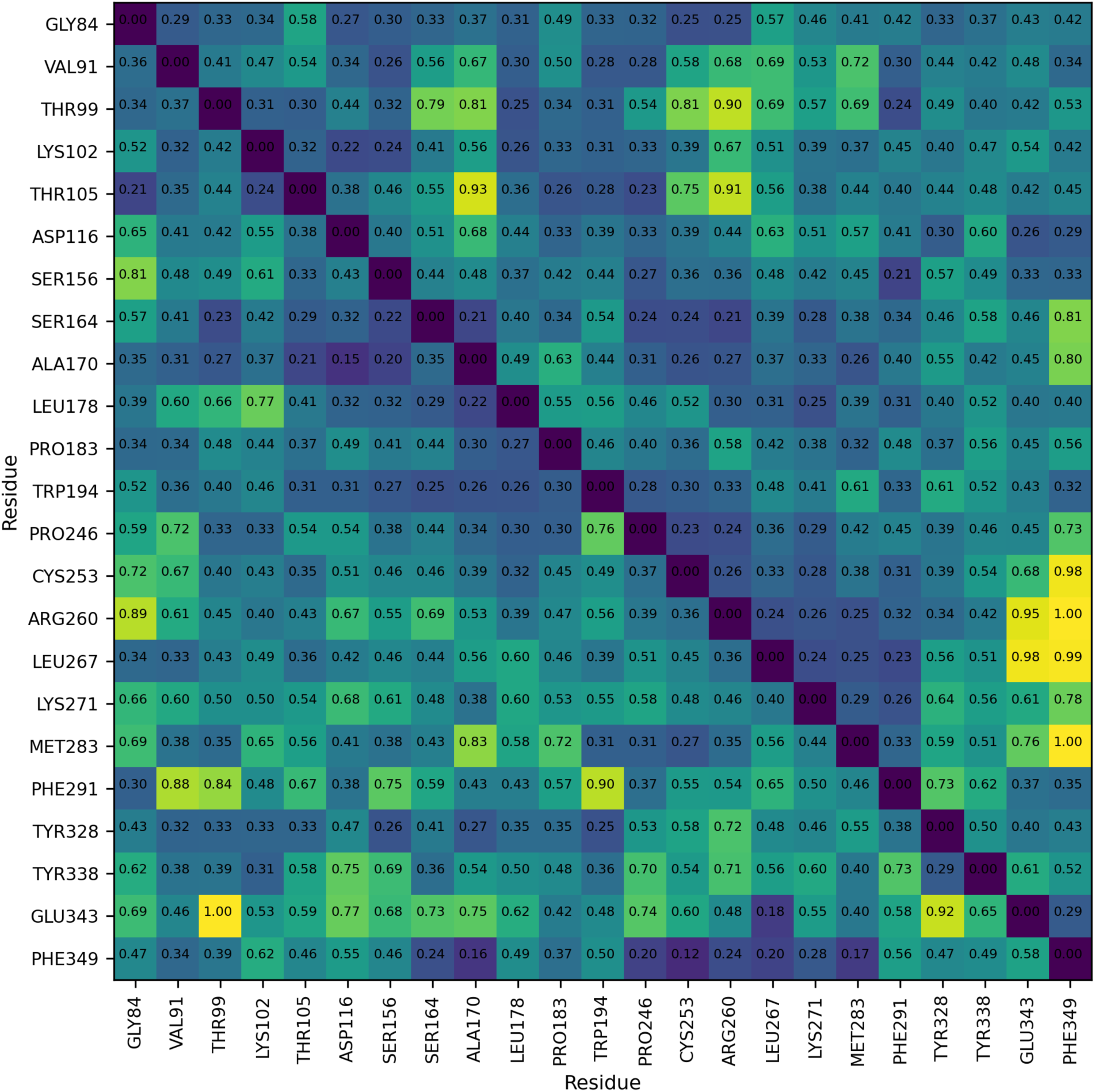
Efficacy response function scores for pairwise distances between alpha carbons from residues in the middle and intracellular end of each helix, with at least one residue in between, and the middle of each intracellular loop. Scores below the diagonal were calculated with the general activation function (Equation 8). Scores above the diagonal were calculated with the selective activation function (Equation 9). Scores from each category were divided by the largest score in the category.

**Figure S7.**
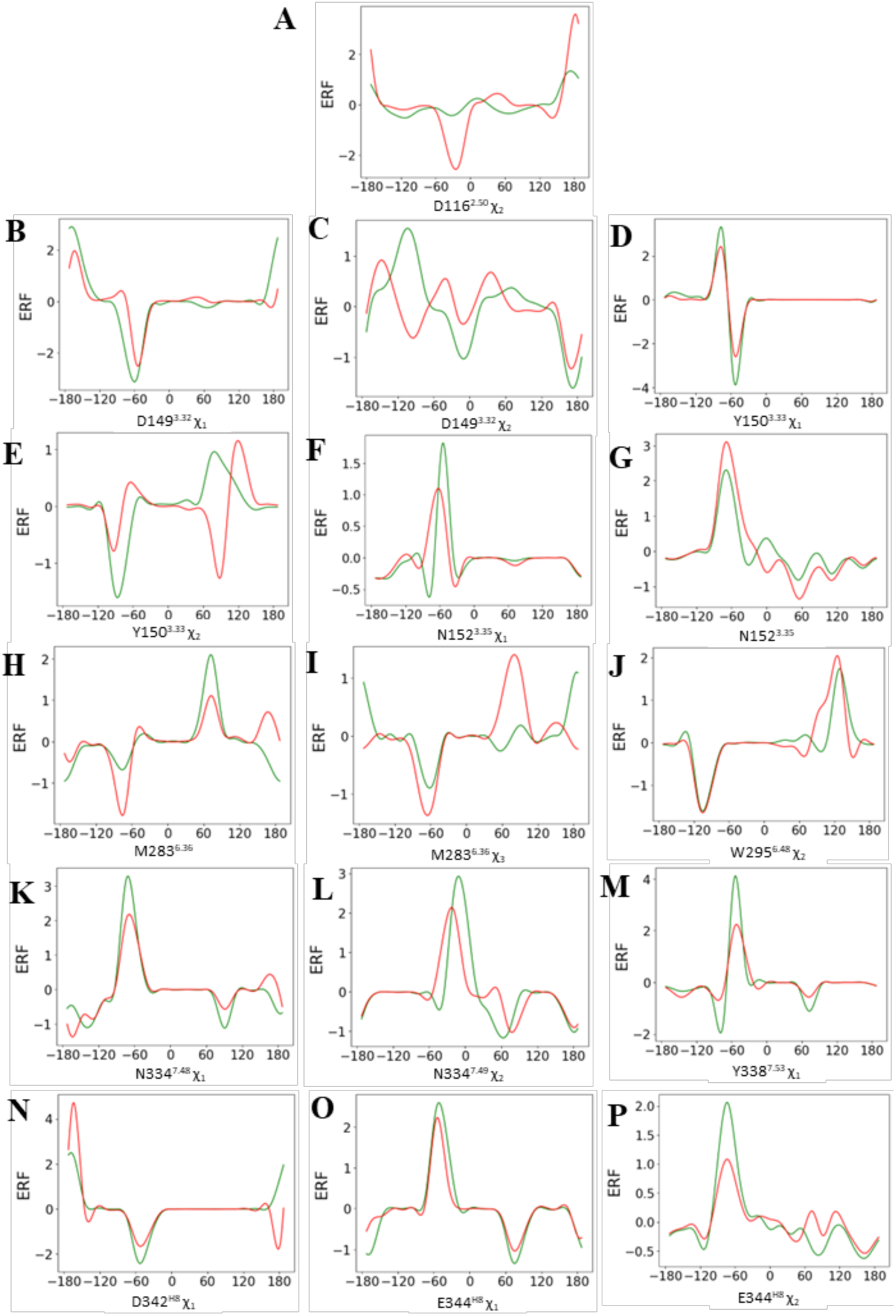
Efficacy response functions for dihedral angles with the largest general activation scores: (A) D116^2.50^ χ_2_ angle, (B) D149^3.32^ χ_1_ angle, (C) D149^3.32^ χ_2_ angle, (D) Y150^3.33^ χ_1_ angle, (E) Y150^3.33^ χ_2_ angle, (F) N152^3.35^ χ_1_ angle, (G) N152^3.35^ χ_2_ angle, (H) M283^6.36^ χ_2_ angle, (I) M283^6.36^ χ_3_ angle, (J) W295^6.48^ χ_2_ angle, (K) N334^7.48^ χ_1_ angle, (L) N334^7.49^ χ_2_ angle, (M) Y338^7.53^ χ_1_ angle, (N) D342^H8^ χ_1_ angl^e,^ (O) E344^H8^ χ_1_ angle, (P) E344^H8^ χ_2_ angle. The green and red curves correspond to the G protein and β-arrestin-2 efficacy response functions (ERF), respectively.

**Figure S8.**
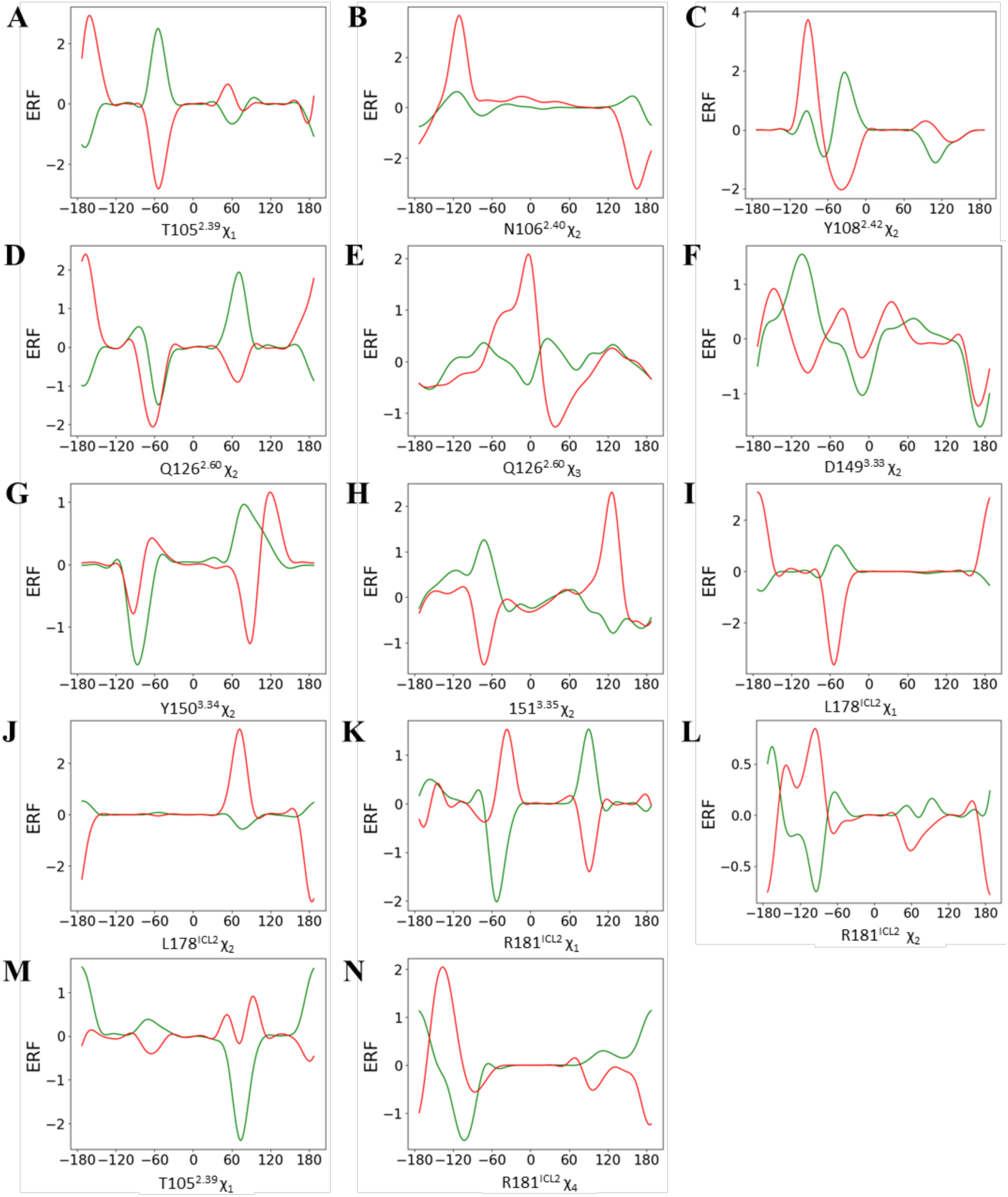
Efficacy response functions for dihedral angles with the largest selective activation scores: (A) T105^2.39^ χ_1_ angle, (B) N106^2.40^ χ_2_ angle, (C) Y108^2.42^ χ_2_ angle, (D) Q126^2.60^ χ_2_ angle, (E) Q126^2.60^ χ_3_ angle, (F) D149^3.32^ χ_2_ angle, (G) Y150^3.33^ χ_2_ angle, (H) 151^3.35^ χ_2_ angle, (I) L178^ICL2^ χ_1_ angle, (J) L178^ICL2^ χ_2_ angle, (K) R181^ICL2^ χ1 angle, (L) R181ICL2 χ_2_ angle, (M) R181^ICL2^ χ_3_ angle, (N) R181^ICL2^ χ_4_ angle. The green and red curves correspond to the G protein and β-arrestin-2 efficacy response functions (ERF), respectively.

**Figure S9.**
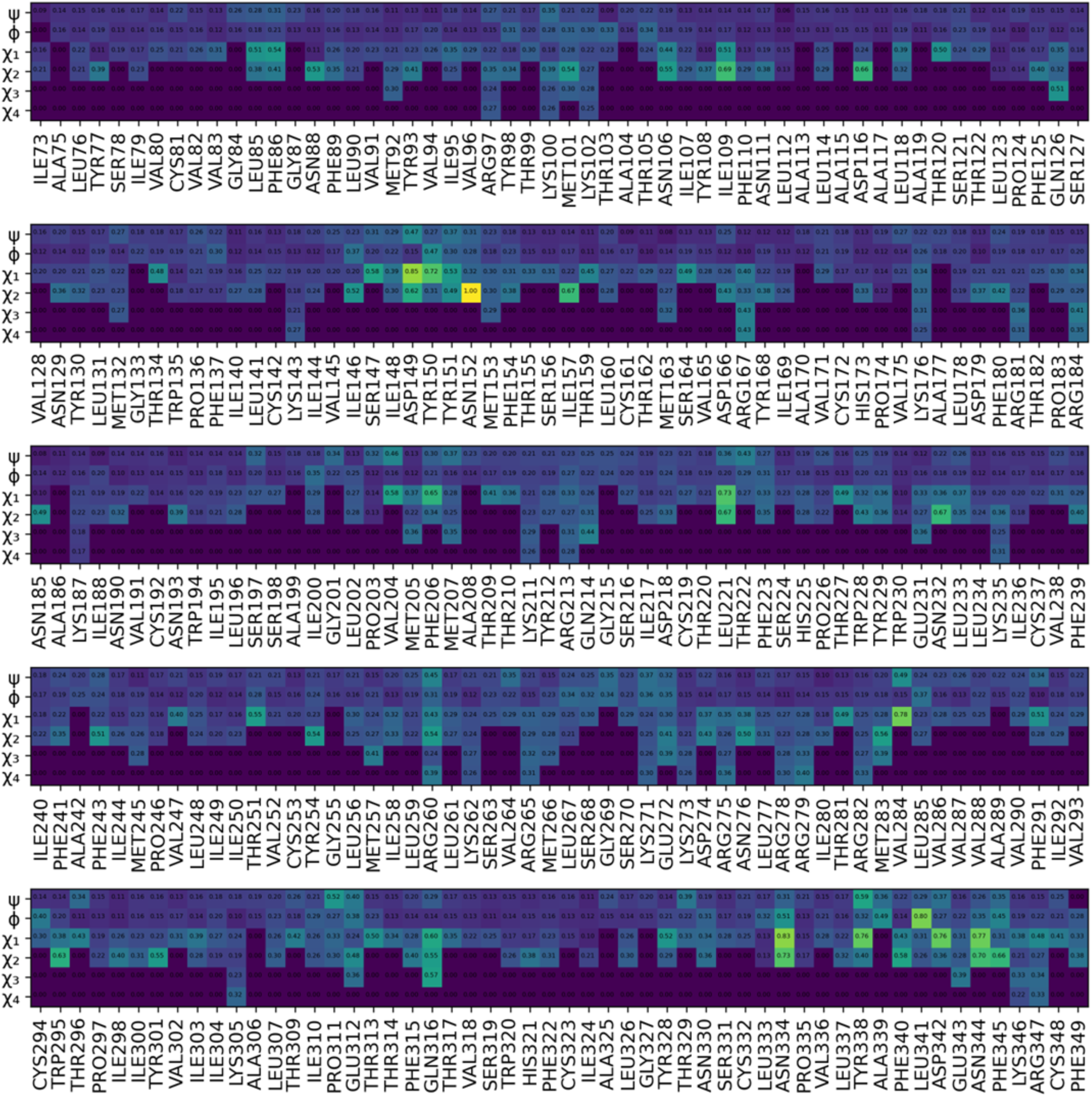
General activation function scores for amino acid dihedral angles.

**Figure S10.**
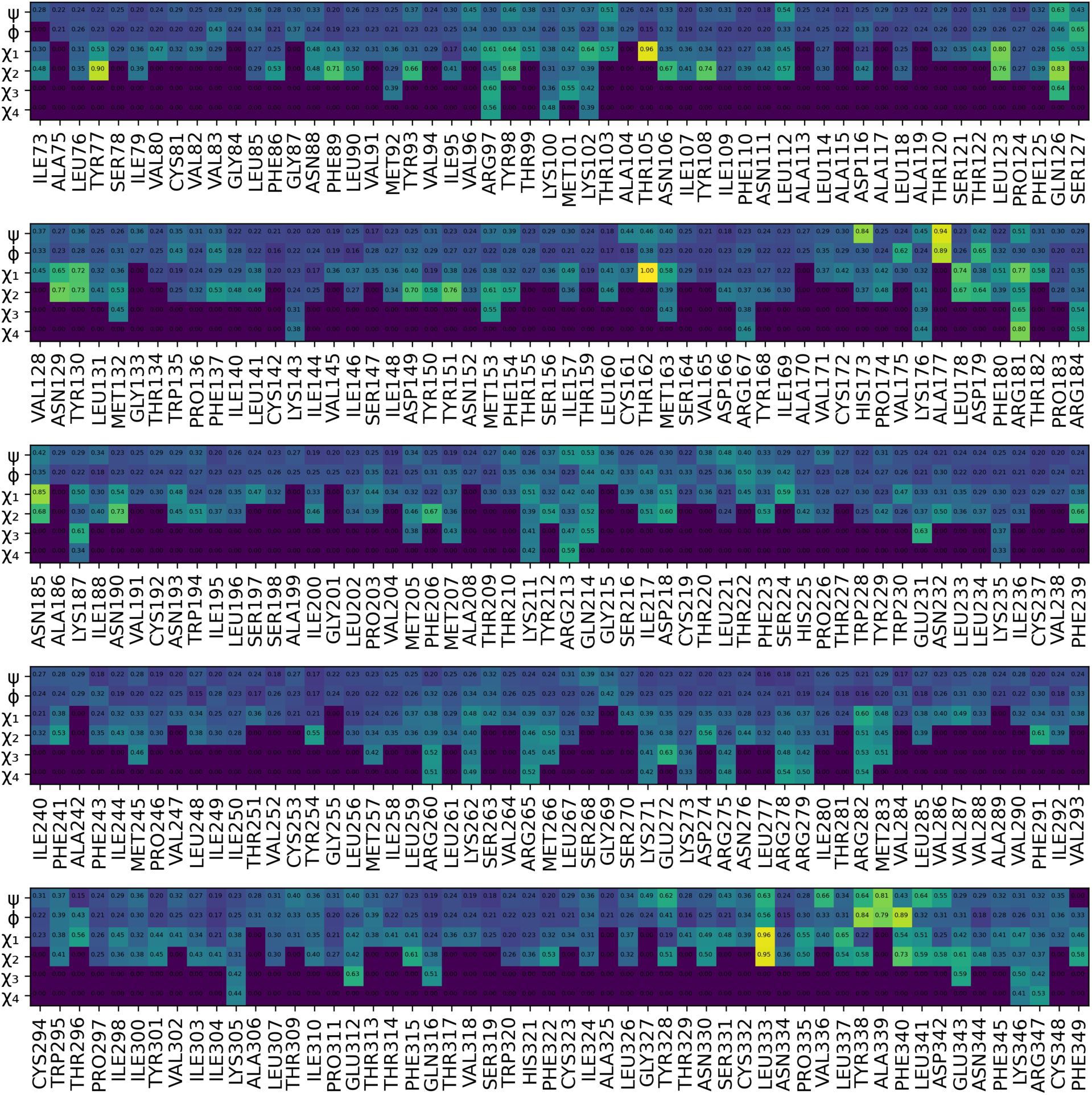
Selective activation function scores for amino acid dihedral angles.

**Figure S11.**
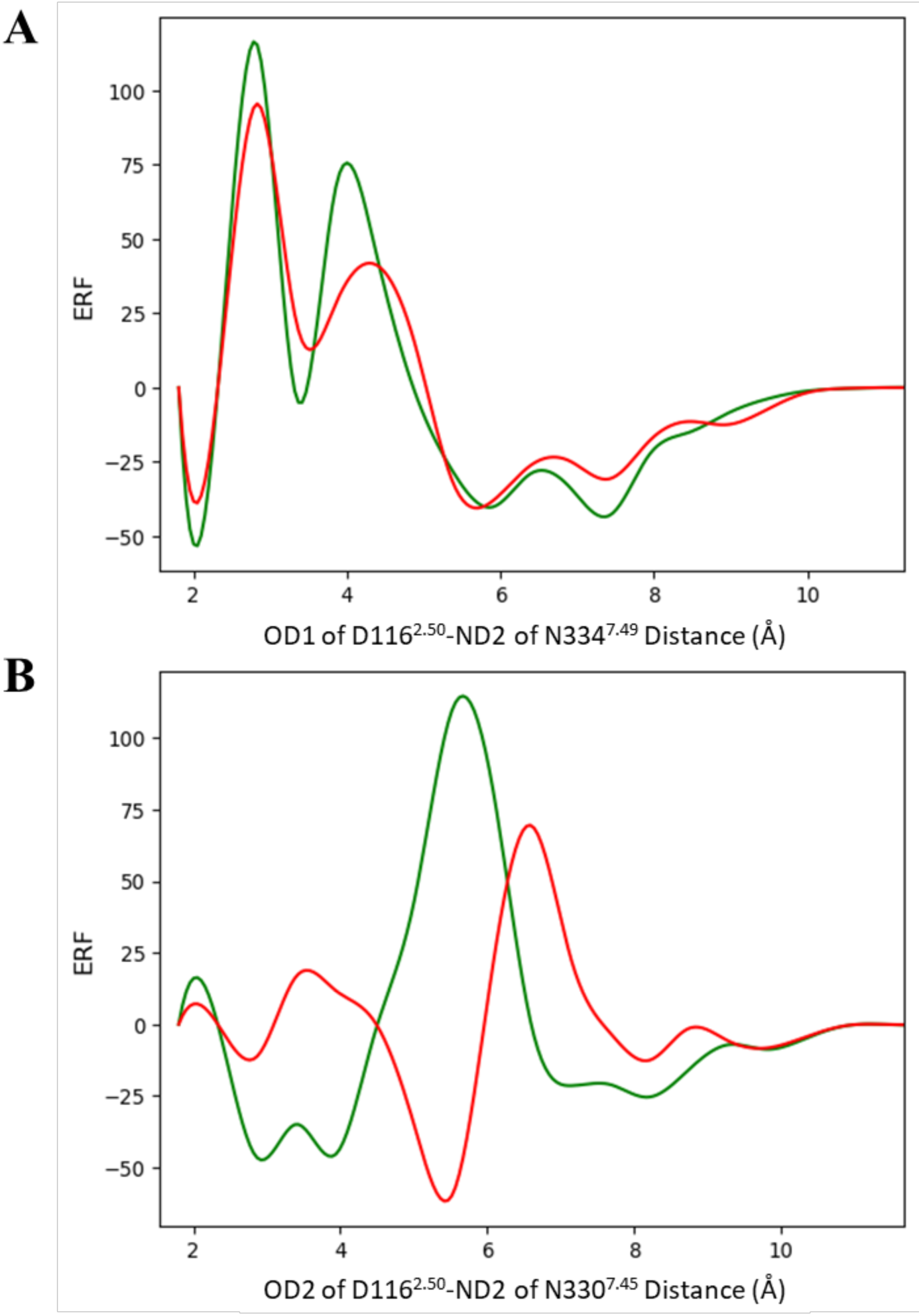
Efficacy response functions of the distance between sodium pocket polar atoms: (A) OD1 of D116^2.50^ and ND2 of N334^7.49^ and (B) OD2 of D116^2.50^ and ND2 of N330^7.45^. The green and red curves correspond to the G protein and β-arrestin-2 efficacy response functions (ERF), respectively.

**Figure S12.**
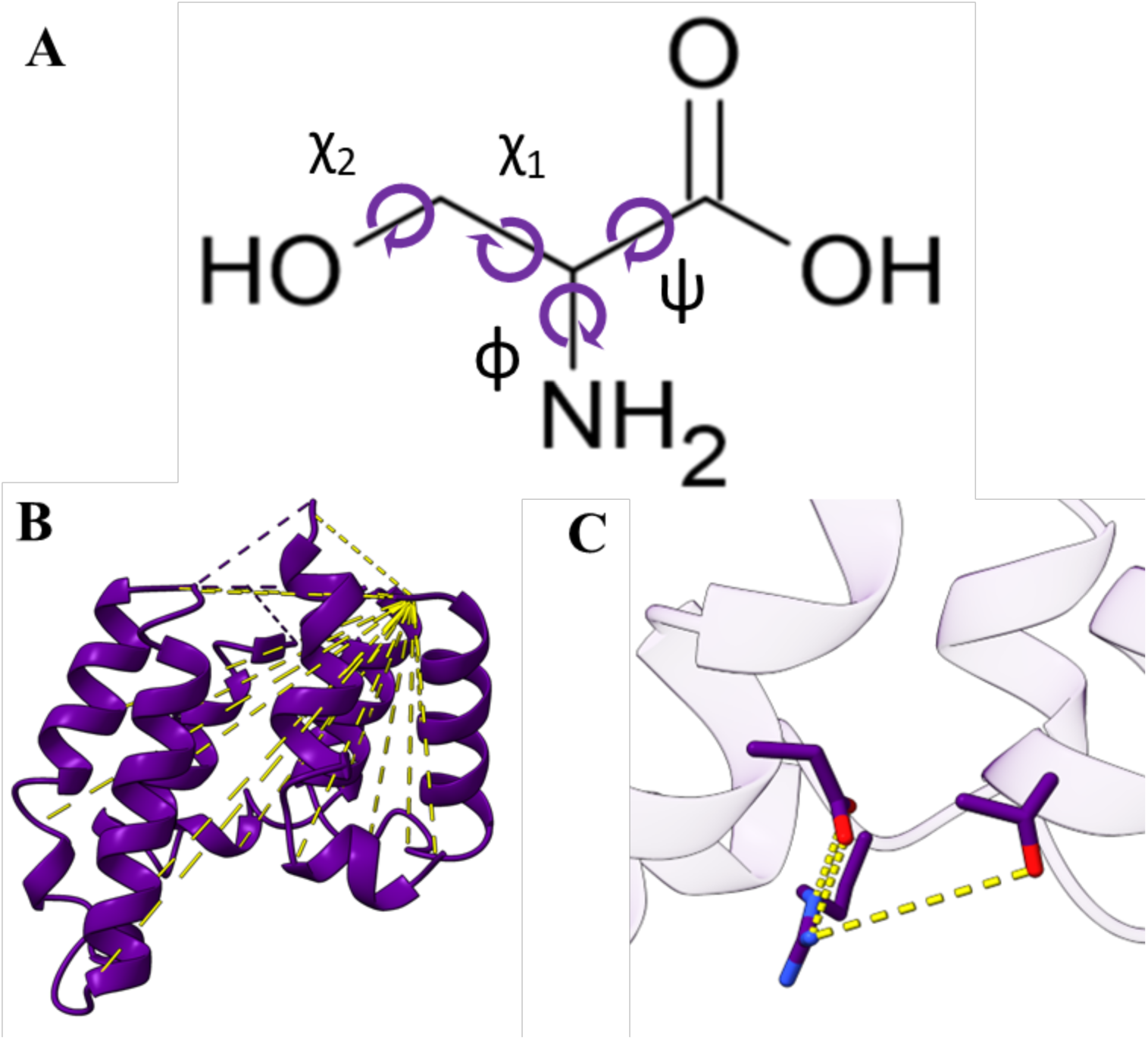
Representations of features used to discretize configurations into conformations: A) dihedral angles, B) alpha carbon distances, and C) distances between polar atoms in the intracellular region of the μOR.

**Table S1:**
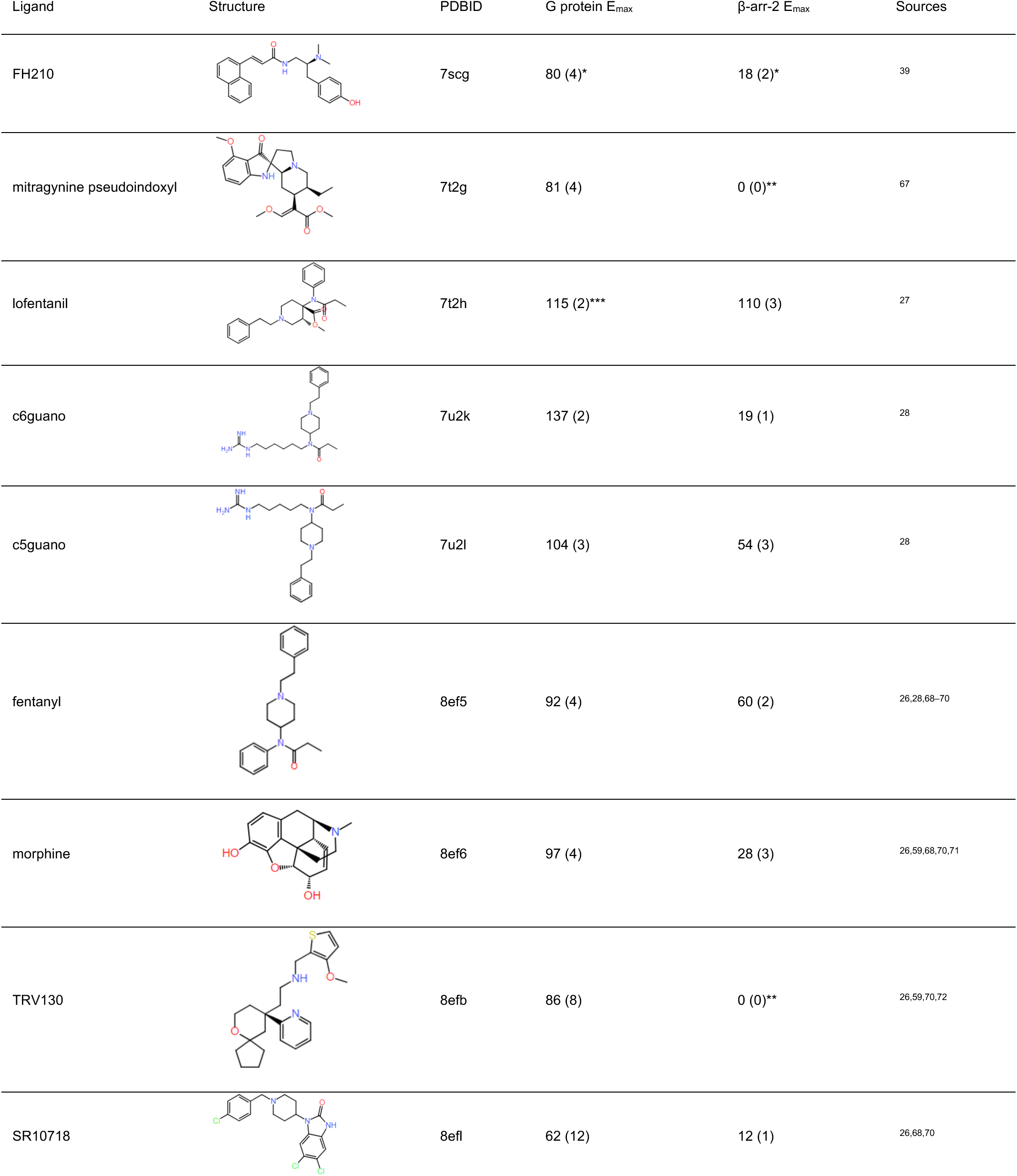
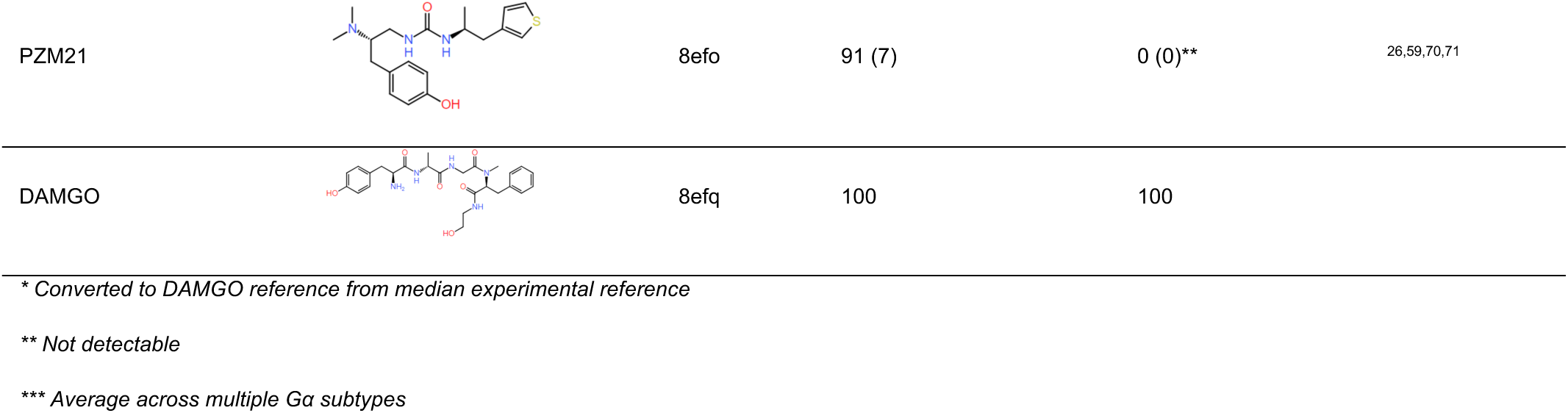
Summary of structures and efficacy data used for MDS and model training. Emax reported as median Emax (median STD)

| Ligand | Structure | PDBID | G protein E<sub>max</sub> | β-arr-2 E<sub>max</sub> | Sources |
| --- | --- | --- | --- | --- | --- |
| FH210 |  | 7scg | 80 (4)* | 18 (2)* | 39 |
| mitragynine pseudoindoxyl |  | 7t2g | 81 (4) | 0 (0)** | 67 |
| lofentanil |  | 7t2h | 115 (2)*** | 110 (3) | 27 |
| c6guano |  | 7u2k | 137 (2) | 19 (1) | 28 |
| c5guano |  | 7u2l | 104 (3) | 54 (3) | 28 |
| fentanyl |  | 8ef5 | 92 (4) | 60 (2) | 26,28,68–70 |
| morphine |  | 8ef6 | 97 (4) | 28 (3) | 26,59,68,70,71 |
| TRV130 |  | 8efb | 86 (8) | 0 (0)** | 26,59,70,72 |
| SR10718 |  | 8efl | 62 (12) | 12 (1) | 26,68,70 |
| PZM21 |  | 8efo | 91 (7) | 0 (0)** | 26,59,70,71 |
| DAMGO |  | 8efq | 100 | 100 |  |

\* Converted to DAMGO reference from median experimental reference
\*\* Not detectable
\*\*\* Average across multiple Ga subtypes

**Table S2:**
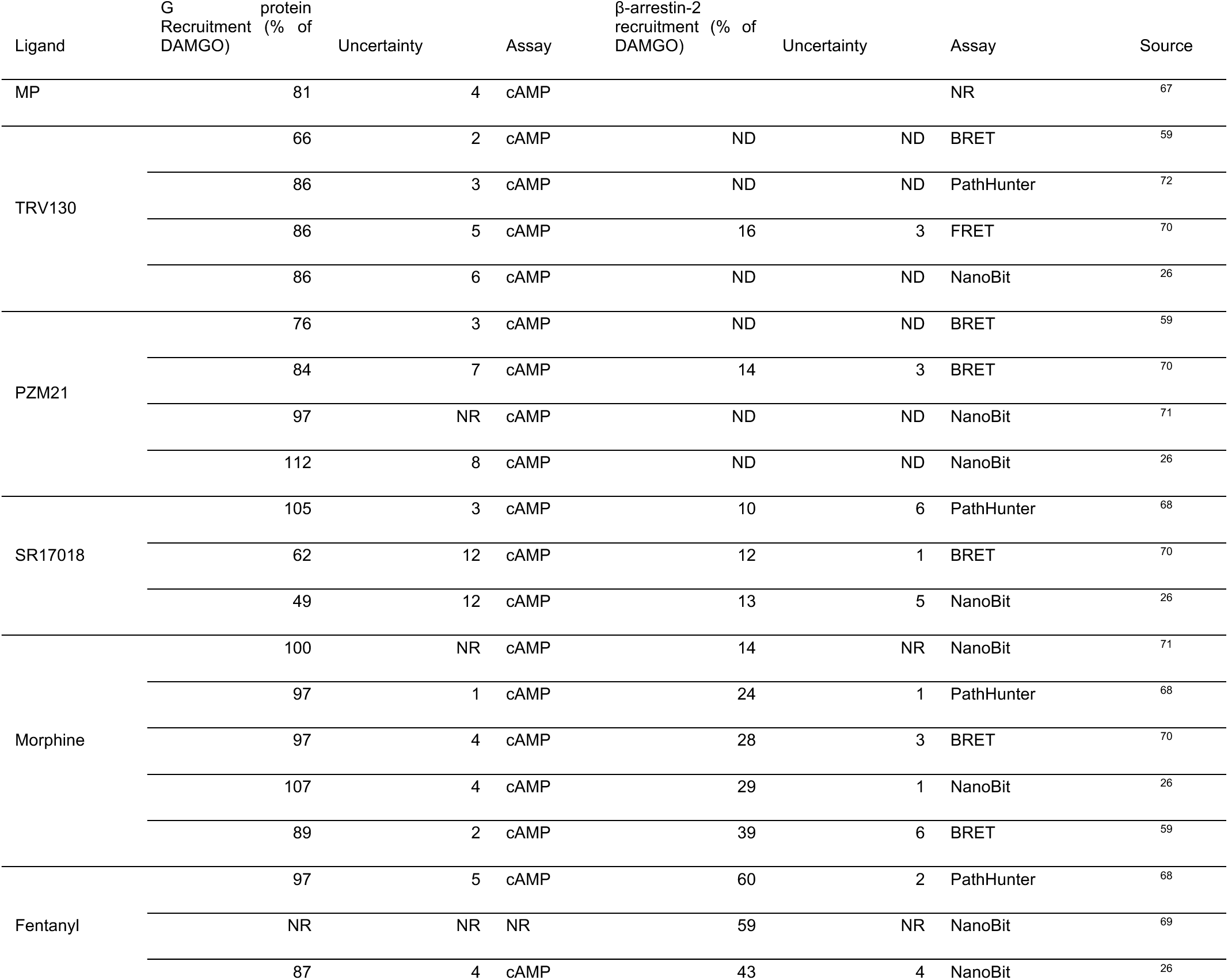
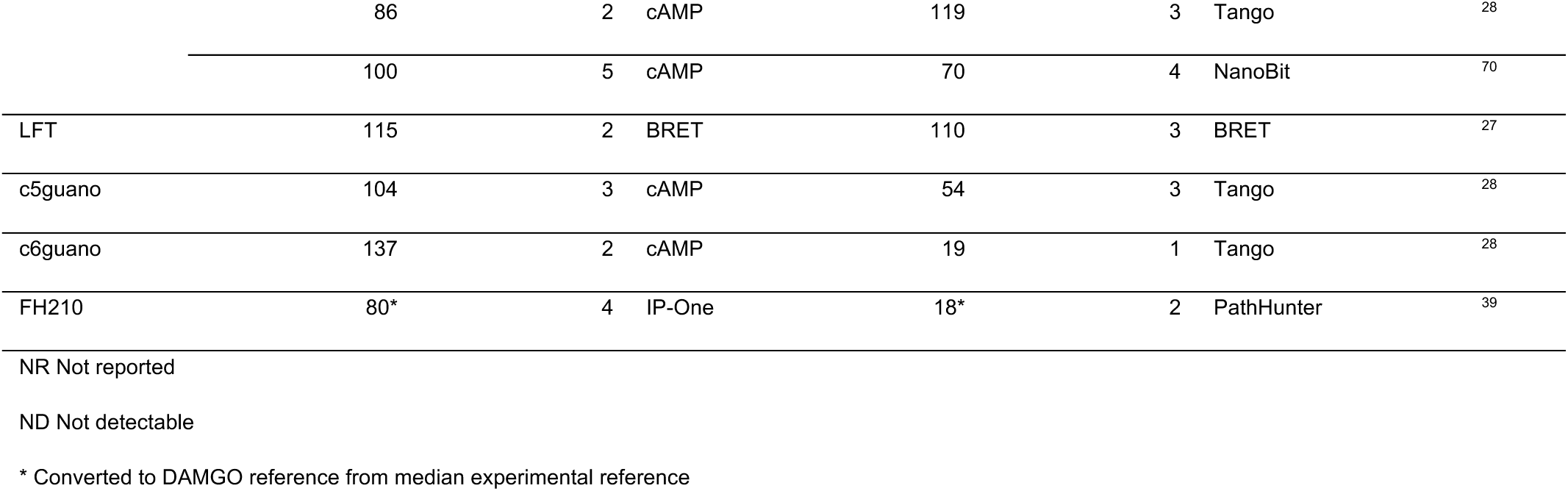
Experimental cAMP and βarr2 efficacies used to compute the median experimental efficacies for model training.

## REFERENCES

(1) Smith, J. S.; Lefkowitz, R. J.; Rajagopal, S. Biased Signalling: From Simple Switches to Allosteric Microprocessors. Nat. Rev. Drug Discov. 2018, 17 (4), 243–260. 10.1038/nrd.2017.229.

(2) Hauser, A. S.; Attwood, M. M.; Rask-Andersen, M.; Schiöth, H. B.; Gloriam, D. E. Trends in GPCR Drug Discovery: New Agents, Targets and Indications. Nat. Rev. Drug Discov. 2017, 16 (12), 829–842. 10.1038/nrd.2017.178.

(3) Whalen, E. J.; Rajagopal, S.; Lefkowitz, R. J. Therapeutic Potential of β-Arrestin- and G Protein-Biased Agonists. Trends Mol. Med. 2011, 17 (3), 126–139. 10.1016/j.molmed.2010.11.004.

(4) Tan, L.; Yan, W.; McCorvy, J. D.; Cheng, J. Biased Ligands of G Protein-Coupled Receptors (GPCRs): Structure–Functional Selectivity Relationships (SFSRs) and Therapeutic Potential. J. Med. Chem. 2018, 61 (22), 9841–9878. 10.1021/acs.jmedchem.8b00435.

(5) Seyedabadi, M.; Ghahremani, M. H.; Albert, P. R. Biased Signaling of G Protein Coupled Receptors (GPCRs): Molecular Determinants of GPCR/Transducer Selectivity and Therapeutic Potential. Pharmacol. Ther. 2019, 200, 148–178. 10.1016/j.pharmthera.2019.05.006.

(6) Faouzi, A.; Varga, B. R.; Majumdar, S. Biased Opioid Ligands. Molecules 2020, 25 (18), 4257. 10.3390/molecules25184257.

(7) De Neve, J.; Barlow, T. M. A.; Tourwé, D.; Bihel, F.; Simonin, F.; Ballet, S. Comprehensive Overview of Biased Pharmacology at the Opioid Receptors: Biased Ligands and Bias Factors. RSC Med. Chem. 2021, 12 (6), 828–870. 10.1039/D1MD00041A.

(8) Eiger, D. S.; Pham, U.; Gardner, J.; Hicks, C.; Rajagopal, S. GPCR Systems Pharmacology: A Different Perspective on the Development of Biased Therapeutics. Am. J. Physiol.-Cell Physiol. 2022, 322 (5), C887–C895. 10.1152/ajpcell.00449.2021.

(9) Kelly, E.; Sutcliffe, K.; Cavallo, D.; Ramos-Gonzalez, N.; Alhosan, N.; Henderson, G. The Anomalous Pharmacology of Fentanyl. Br. J. Pharmacol. 2023, 180 (7), 797–812. 10.1111/bph.15573.

(10) Stanley, T. H. The History and Development of the Fentanyl Series. J. Pain Symptom Manage. 1992, 7 (3), S3–S7. 10.1016/0885-3924(92)90047-L.

(11) Leysen, J. E.; Gommeren, W.; Niemegeers, C. J. E. [3H]Sufentanil, a Superior Ligand for μ-Opiate Receptors: Binding Properties and Regional Distribution in Rat Brain and Spinal Cord. Eur. J. Pharmacol. 1983, 87 (2–3), 209–225. 10.1016/0014-2999(83)90331-X.

(12) Spencer, M. R. Drug Overdose Deaths in the United States, 2001–202. 2022, No. 457.

(13) Weis, W. I.; Kobilka, B. K. The Molecular Basis of G Protein–Coupled Receptor Activation. Annu. Rev. Biochem. 2018, 87 (1), 897–919. 10.1146/annurev-biochem-060614-033910.

(14) Wingler, L. M.; Lefkowitz, R. J. Conformational Basis of G Protein-Coupled Receptor Signaling Versatility. Trends Cell Biol. 2020, 30 (9), 736–747. 10.1016/j.tcb.2020.06.002.

(15) Wingler, L. M.; Elgeti, M.; Hilger, D.; Latorraca, N. R.; Lerch, M. T.; Staus, D. P.; Dror, R. O.; Kobilka, B. K.; Hubbell, W. L.; Lefkowitz, R. J. Angiotensin Analogs with Divergent Bias Stabilize Distinct Receptor Conformations. Cell 2019, 176 (3), 468–478.e11. 10.1016/j.cell.2018.12.005.

(16) Lamichhane, R.; Liu, J. J.; White, K. L.; Katritch, V.; Stevens, R. C.; Wüthrich, K.; Millar, D. P. Biased Signaling of the G-Protein-Coupled Receptor β2AR Is Governed by Conformational Exchange Kinetics. Structure 2020, 28 (3), 371–377.e3. 10.1016/j.str.2020.01.001.

(17) Imai, S.; Yokomizo, T.; Kofuku, Y.; Shiraishi, Y.; Ueda, T.; Shimada, I. Structural Equilibrium Underlying Ligand-Dependent Activation of Β2-Adrenoreceptor. Nat. Chem. Biol. 2020, 16 (4), 430–439. 10.1038/s41589-019-0457-5.

(18) Cong et. al., X. Molecular Insights into the Biased Signaling Mechanism of MOR. Mol. Cell 2021, 81 (20), 4165–4175. 10.1016/j.molcel.2021.07.033.

(19) Che, T.; Roth, B. L. Molecular Basis of Opioid Receptor Signaling. Cell 2023, 186 (24), 5203–5219. 10.1016/j.cell.2023.10.029.

(20) Li, Z.; Liu, J.; Dong, F.; Chang, N.; Huang, R.; Xia, M.; Patterson, T. A.; Hong, H. Three-Dimensional Structural Insights Have Revealed the Distinct Binding Interactions of Agonists, Partial Agonists, and Antagonists with the µ Opioid Receptor. Int. J. Mol. Sci. 2023, 24 (8), 7042. 10.3390/ijms24087042.

(21) Shen, S.; Zhao, C.; Wu, C.; Sun, S.; Li, Z.; Yan, W.; Shao, Z. Allosteric Modulation of G Protein-Coupled Receptor Signaling. Front. Endocrinol. 2023, 14, 1137604. 10.3389/fendo.2023.1137604.

(22) Claff, T.; Yu, J.; Blais, V.; Patel, N.; Martin, C.; Wu, L.; Han, G. W.; Holleran, B. J.; Van Der Poorten, O.; White, K. L.; Hanson, M. A.; Sarret, P.; Gendron, L.; Cherezov, V.; Katritch, V.; Ballet, S.; Liu, Z.-J.; Müller, C. E.; Stevens, R. C. Elucidating the Active δ-Opioid Receptor Crystal Structure with Peptide and Small-Molecule Agonists. Sci. Adv. 2019, 5 (11), eaax9115. 10.1126/sciadv.aax9115.

(23) Zhang, D.; Zhao, Q.; Wu, B. Structural Studies of G Protein-Coupled Receptors. Mol. Cells 2015, 38 (10), 836–842. 10.14348/molcells.2015.0263.

(24) Wingler, L. M.; Skiba, M. A.; McMahon, C.; Staus, D. P.; Kleinhenz, A. L. W.; Suomivuori, C.-M.; Latorraca, N. R.; Dror, R. O.; Lefkowitz, R. J.; Kruse, A. C. Angiotensin and Biased Analogs Induce Structurally Distinct Active Conformations within a GPCR. Science 2020, 367 (6480), 888–892. 10.1126/science.aay9813.

(25) Piekielna-Ciesielska, J.; Artali, R.; Azzam, A. A. H.; Lambert, D. G.; Kluczyk, A.; Gentilucci, L.; Janecka, A. Pharmacological Characterization of Μ-Opioid Receptor Agonists with Biased G Protein or β-Arrestin Signaling, and Computational Study of Conformational Changes during Receptor Activation. Molecules 2020, 26 (1), 13. 10.3390/molecules26010013.

(26) Zhuang, Y.; Wang, Y.; He, B.; He, X.; Zhou, X. E.; Guo, S.; Rao, Q.; Yang, J.; Liu, J.; Zhou, Q.; Wang, X.; Liu, M.; Liu, W.; Jiang, X.; Yang, D.; Jiang, H.; Shen, J.; Melcher, K.; Chen, H.; Jiang, Y.; Cheng, X.; Wang, M.-W.; Xie, X.; Xu, H. E. Molecular Recognition of Morphine and Fentanyl by the Human μ-Opioid Receptor. Cell 2022, 185 (23), 4361–4375.e19. 10.1016/j.cell.2022.09.041.

(27) Qu, Q.; Huang, W.; Aydin, D.; Paggi, J. M.; Seven, A. B.; Wang, H.; Chakraborty, S.; Che, T.; DiBerto, J. F.; Robertson, M. J.; Inoue, A.; Suomivuori, C.-M.; Roth, B. L.; Majumdar, S.; Dror, R. O.; Kobilka, B. K.; Skiniotis, G. Insights into Distinct Signaling Profiles of the µOR Activated by Diverse Agonists. Nat. Chem. Biol. 2023, 19 (4), 423–430. 10.1038/s41589-022-01208-y.

(28) Faouzi, A.; Wang, H.; Zaidi, S. A.; DiBerto, J. F.; Che, T.; Qu, Q.; Robertson, M. J.; Madasu, M. K.; El Daibani, A.; Varga, B. R.; Zhang, T.; Ruiz, C.; Liu, S.; Xu, J.; Appourchaux, K.; Slocum, S. T.; Eans, S. O.; Cameron, M. D.; Al-Hasani, R.; Pan, Y. X.; Roth, B. L.; McLaughlin, J. P.; Skiniotis, G.; Katritch, V.; Kobilka, B. K.; Majumdar, S. Structure-Based Design of Bitopic Ligands for the µ-Opioid Receptor. Nature 2023, 613 (7945), 767–774. 10.1038/s41586-022-05588-y.

(29) Dolinsky, T. J.; Czodrowski, P.; Li, H.; Nielsen, J. E.; Jensen, J. H.; Klebe, G.; Baker, N. A. PDB2PQR: Expanding and Upgrading Automated Preparation of Biomolecular Structures for Molecular Simulations. Nucleic Acids Res. 2007, 35 (Web Server), W522–W525. 10.1093/nar/gkm276.

(30) Lomize, A. L.; Pogozheva, I. D.; Mosberg, H. I. Anisotropic Solvent Model of the Lipid Bilayer. 2. Energetics of Insertion of Small Molecules, Peptides, and Proteins in Membranes. J. Chem. Inf. Model. 2011, 51 (4), 930–946. 10.1021/ci200020k.

(31) Maier, J. A.; Martinez, C.; Kasavajhala, K.; Wickstrom, L.; Hauser, K. E.; Simmerling, C. ff14SB: Improving the Accuracy of Protein Side Chain and Backbone Parameters from ff99SB. J. Chem. Theory Comput. 2015, 11 (8), 3696–3713. 10.1021/acs.jctc.5b00255.

(32) Izadi, S.; Anandakrishnan, R.; Onufriev, A. V. Building Water Models: A Different Approach. J. Phys. Chem. Lett. 2014, 5 (21), 3863–3871. 10.1021/jz501780a.

(33) Case, D. A.; Aktulga, H. M.; Belfon, K.; Cerutti, D. S.; Cisneros, G. A.; Cruzeiro, V. W. D.; Forouzesh, N.; Giese, T. J.; Götz, A. W.; Gohlke, H.; Izadi, S.; Kasavajhala, K.; Kaymak, M. C.; King, E.; Kurtzman, T.; Lee, T.-S.; Li, P.; Liu, J.; Luchko, T.; Luo, R.; Manathunga, M.; Machado, M. R.; Nguyen, H. M.; O’Hearn, K. A.; Onufriev, A. V.; Pan, F.; Pantano, S.; Qi, R.; Rahnamoun, A.; Risheh, A.; Schott-Verdugo, S.; Shajan, A.; Swails, J.; Wang, J.; Wei, H.; Wu, X.; Wu, Y.; Zhang, S.; Zhao, S.; Zhu, Q.; Cheatham, T. E.; Roe, D. R.; Roitberg, A.; Simmerling, C.; York, D. M.; Nagan, M. C.; Merz, K. M. AmberTools. J. Chem. Inf. Model. 2023, 63 (20), 6183–6191. 10.1021/acs.jcim.3c01153.

(34) Eastman, P.; Swails, J.; Chodera, J. D.; McGibbon, R. T.; Zhao, Y.; Beauchamp, K. A.; Wang, L.-P.; Simmonett, A. C.; Harrigan, M. P.; Stern, C. D.; Wiewiora, R. P.; Brooks, B. R.; Pande, V. S. OpenMM 7: Rapid Development of High Performance Algorithms for Molecular Dynamics. PLOS Comput. Biol. 2017, 13 (7), e1005659. 10.1371/journal.pcbi.1005659.

(35) Boerner, T. J.; Deems, S.; Furlani, T. R.; Knuth, S. L.; Towns, J. ACCESS: Advancing Innovation: NSF’s Advanced Cyberinfrastructure Coordination Ecosystem: Services & Support. In Practice and Experience in Advanced Research Computing; ACM: Portland OR USA, 2023; pp 173–176. 10.1145/3569951.3597559.

(36) Zhang, R.; Xie, X. Tools for GPCR Drug Discovery. Acta Pharmacol. Sin. 2012, 33 (3), 372–384. 10.1038/aps.2011.173.

(37) Pedregosa, F.; Varoquaux, G.; Gramfort, A.; Michel, V.; Thirion, B.; Grisel, O.; Blondel, M.; Prettenhofer, P.; Weiss, R.; Dubourg, V.; Vanderplas, J.; Passos, A.; Cournapeau, D.; Brucher, M.; Perrot, M.; Duchesnay, E. Scikit-Learn: Machine Learning in {P}ython. J. Mach. Learn. Res. 2011, 12, 2825–2830.

(38) Ballesteros, J. A.; Weinstein, H. [19] Integrated Methods for the Construction of Three-Dimensional Models and Computational Probing of Structure-Function Relations in G Protein-Coupled Receptors. In Methods in Neurosciences; Sealfon, S. C., Ed.; Receptor Molecular Biology; Academic Press, 1995; Vol. 25, pp 366–428. 10.1016/S1043-9471(05)80049-7.

(39) Wang, H.; Hetzer, F.; Huang, W.; Qu, Q.; Meyerowitz, J.; Kaindl, J.; Hübner, H.; Skiniotis, G.; Kobilka, B. K.; Gmeiner, P. Structure-Based Evolution of G Protein-Biased μ-Opioid Receptor Agonists. Angew. Chem. Int. Ed. 2022, 61 (26), e202200269. 10.1002/anie.202200269.

(40) Virtanen, P.; Gommers, R.; Oliphant, T. E.; Haberland, M.; Reddy, T.; Cournapeau, D.; Burovski, E.; Peterson, P.; Weckesser, W.; Bright, J.; van der Walt, S. J.; Brett, M.; Wilson, J.; Millman, K. J.; Mayorov, N.; Nelson, A. R. J.; Jones, E.; Kern, R.; Larson, E.; Carey, C. J.; Polat, İ.; Feng, Y.; Moore, E. W.; VanderPlas, J.; Laxalde, D.; Perktold, J.; Cimrman, R.; Henriksen, I.; Quintero, E. A.; Harris, C. R.; Archibald, A. M.; Ribeiro, A. H.; Pedregosa, F.; van Mulbregt, P.; SciPy 1.0 Contributors; Vijaykumar, A.; Bardelli, A. P.; Rothberg, A.; Hilboll, A.; Kloeckner, A.; Scopatz, A.; Lee, A.; Rokem, A.; Woods, C. N.; Fulton, C.; Masson, C.; Häggström, C.; Fitzgerald, C.; Nicholson, D. A.; Hagen, D. R.; Pasechnik, D. V.; Olivetti, E.; Martin, E.; Wieser, E.; Silva, F.; Lenders, F.; Wilhelm, F.; Young, G.; Price, G. A.; Ingold, G.-L.; Allen, G. E.; Lee, G. R.; Audren, H.; Probst, I.; Dietrich, J. P.; Silterra, J.; Webber, J. T.; Slavič, J.; Nothman, J.; Buchner, J.; Kulick, J.; Schönberger, J. L.; de Miranda Cardoso, J. V.; Reimer, J.; Harrington, J.; Rodríguez, J. L. C.; Nunez-Iglesias, J.; Kuczynski, J.; Tritz, K.; Thoma, M.; Newville, M.; Kümmerer, M.; Bolingbroke, M.; Tartre, M.; Pak, M.; Smith, N. J.; Nowaczyk, N.; Shebanov, N.; Pavlyk, O.; Brodtkorb, P. A.; Lee, P.; McGibbon, R. T.; Feldbauer, R.; Lewis, S.; Tygier, S.; Sievert, S.; Vigna, S.; Peterson, S.; More, S.; Pudlik, T.; Oshima, T.; Pingel, T. J.; Robitaille, T. P.; Spura, T.; Jones, T. R.; Cera, T.; Leslie, T.; Zito, T.; Krauss, T.; Upadhyay, U.; Halchenko, Y. O.; Vázquez-Baeza, Y. SciPy 1.0: Fundamental Algorithms for Scientific Computing in Python. Nat. Methods 2020, 17 (3), 261–272. 10.1038/s41592-019-0686-2.

(41) van der Walt, S.; Colbert, S. C.; Varoquaux, G. The NumPy Array: A Structure for Efficient Numerical Computation. Comput. Sci. Eng. 2011, 13 (2), 22–30. 10.1109/MCSE.2011.37.

(42) Miao, Y.; McCammon, J. A. Graded Activation and Free Energy Landscapes of a Muscarinic G-Protein–Coupled Receptor. Proc. Natl. Acad. Sci. 2016, 113 (43), 12162–12167. 10.1073/pnas.1614538113.

(43) Suomivuori, C.-M.; Latorraca, N. R.; Wingler, L. M.; Eismann, S.; King, M. C.; Kleinhenz, A. L. W.; Skiba, M. A.; Staus, D. P.; Kruse, A. C.; Lefkowitz, R. J.; Dror, R. O. Molecular Mechanism of Biased Signaling in a Prototypical G Protein–Coupled Receptor. Science 2020, 367, 881–887. 10.1126/science.aaz0326.

(44) Panel, N.; Vo, D. D.; Kahlous, N. A.; Hübner, H.; Tiedt, S.; Matricon, P.; Pacalon, J.; Fleetwood, O.; Kampen, S.; Luttens, A.; Delemotte, L.; Kihlberg, J.; Gmeiner, P.; Carlsson, J. Design of Drug Efficacy Guided by Free Energy Simulations of the β _2_ -Adrenoceptor. Angew. Chem. Int. Ed. 2023, 62 (22), e202218959. 10.1002/anie.202218959.

(45) Vögele, M.; Zhang, B. W.; Kaindl, J.; Wang, L. Is the Functional Response of a Receptor Determined by the Thermodynamics of Ligand Binding? J. Chem. Theory Comput. 2023, 19 (22), 8414–8422. 10.1021/acs.jctc.3c00899.

(46) Dror, R. O.; Arlow, D. H.; Maragakis, P.; Mildorf, T. J.; Pan, A. C.; Xu, H.; Borhani, D. W.; Shaw, D. E. Activation Mechanism of the *β*_2_ -Adrenergic Receptor. Proc. Natl. Acad. Sci. 2011, 108 (46), 18684–18689. 10.1073/pnas.1110499108.

(47) Black, J. W.; Leff, P. Operational Models of Pharmacological Agonism. Proc. R. Soc. Lond. B Biol. Sci. 1983, 220 (1219), 141–162.

(48) Olsen, R. H. J.; English, J. G. Advancements in G Protein-coupled Receptor Biosensors to Study GPCR-G Protein Coupling. Br. J. Pharmacol. 2023, 180 (11), 1433–1443. 10.1111/bph.15962.

(49) Lane, J. R.; May, L. T.; Parton, R. G.; Sexton, P. M.; Christopoulos, A. A Kinetic View of GPCR Allostery and Biased Agonism. Nat. Chem. Biol. 2017, 13 (9), 929–937. 10.1038/nchembio.2431.

(50) Sykes, D. A.; Dowling, M. R.; Charlton, S. J. Exploring the Mechanism of Agonist Efficacy: A Relationship between Efficacy and Agonist Dissociation Rate at the Muscarinic M _3_ Receptor. Mol. Pharmacol. 2009, 76 (3), 543–551. 10.1124/mol.108.054452.

(51) Guo, D.; Mulder-Krieger, T.; IJzerman, A. P.; Heitman, L. H. Functional Efficacy of Adenosine A _2A_ Receptor Agonists Is Positively Correlated to Their Receptor Residence Time. Br. J. Pharmacol. 2012, 166 (6), 1846–1859. 10.1111/j.1476-5381.2012.01897.x.

(52) Louvel, J.; Guo, D.; Soethoudt, M.; Mocking, T. A. M.; Lenselink, E. B.; Mulder-Krieger, T.; Heitman, L. H.; IJzerman, A. P. Structure-Kinetics Relationships of Capadenoson Derivatives as Adenosine A1 Receptor Agonists. Eur. J. Med. Chem. 2015, 101, 681–691. 10.1016/j.ejmech.2015.07.023.

(53) Klein Herenbrink, C.; Sykes, D. A.; Donthamsetti, P.; Canals, M.; Coudrat, T.; Shonberg, J.; Scammells, P. J.; Capuano, B.; Sexton, P. M.; Charlton, S. J.; Javitch, J. A.; Christopoulos, A.; Lane, J. R. The Role of Kinetic Context in Apparent Biased Agonism at GPCRs. Nat. Commun. 2016, 7 (1), 10842. 10.1038/ncomms10842.

(54) Soethoudt, M.; Hoorens, M. W. H.; Doelman, W.; Martella, A.; Van Der Stelt, M.; Heitman, L. H. Structure-Kinetic Relationship Studies of Cannabinoid CB 2 Receptor Agonists Reveal Substituent-Specific Lipophilic Effects on Residence Time. Biochem. Pharmacol. 2018, 152, 129–142. 10.1016/j.bcp.2018.03.018.

(55) Huang, W.; Manglik, A.; Venkatakrishnan, A. J.; Laeremans, T.; Feinberg, E. N.; Sanborn, A. L.; Kato, H. E.; Livingston, K. E.; Thorsen, T. S.; Kling, R. C.; Granier, S.; Gmeiner, P.; Husbands, S. M.; Traynor, J. R.; Weis, W. I.; Steyaert, J.; Dror, R. O.; Kobilka, B. K. Structural Insights into Μ-Opioid Receptor Activation. Nature 2015, 524 (7565), 315–321. 10.1038/nature14886.

(56) Sounier, R.; Mas, C.; Steyaert, J.; Laeremans, T.; Manglik, A.; Huang, W.; Kobilka, B. K.; Déméné, H.; Granier, S. Propagation of Conformational Changes during μ-Opioid Receptor Activation. Nature 2015, 524 (7565), 375–378. 10.1038/nature14680.

(57) Mafi, A.; Kim, S.-K.; Goddard, W. A. Mechanism of β-Arrestin Recruitment by the μ-Opioid G Protein-Coupled Receptor. Proc. Natl. Acad. Sci. U. S. A. 2020, 117 (28), 16346–16355. 10.1073/pnas.1918264117.

(58) Kelly, B.; Hollingsworth, S. A.; Blakemore, D. C.; Owen, R. M.; Storer, R. I.; Swain, N. A.; Aydin, D.; Torella, R.; Warmus, J. S.; Dror, R. O. Delineating the Ligand–Receptor Interactions That Lead to Biased Signaling at the μ-Opioid Receptor. J. Chem. Inf. Model. 2021, 61 (7), 3696–3707. 10.1021/acs.jcim.1c00585.

(59) Manglik, A.; Lin, H.; Aryal, D. K.; McCorvy, J. D.; Dengler, D.; Corder, G.; Levit, A.; Kling, R. C.; Bernat, V.; Hübner, H.; Huang, X.-P.; Sassano, M. F.; Giguère, P. M.; Löber, S.; Da Duan; Scherrer, G.; Kobilka, B. K.; Gmeiner, P.; Roth, B. L.; Shoichet, B. K. Structure-Based Discovery of Opioid Analgesics with Reduced Side Effects. Nature 2016, 537 (7619), 185–190. 10.1038/nature19112.

(60) Wang, W.-W.; Ji, S.-Y.; Zhang, W.; Zhang, J.; Cai, C.; Hu, R.; Zang, S.-K.; Miao, L.; Xu, H.; Chen, L.-N.; Yang, Z.; Guo, J.; Qin, J.; Shen, D.-D.; Liang, P.; Zhang, Y.; Zhang, Y. Structure-Based Design of Non-Hypertrophic Apelin Receptor Modulator. Cell 2024, 187 (6), 1460–1475.e20. 10.1016/j.cell.2024.02.004.

(61) Shang, Y.; LeRouzic, V.; Schneider, S.; Bisignano, P.; Pasternak, G. W.; Filizola, M. Mechanistic Insights into the Allosteric Modulation of Opioid Receptors by Sodium Ions. Biochemistry 2014, 53 (31), 5140–5149. 10.1021/bi5006915.

(62) Marmolejo-Valencia, A. F.; Madariaga-Mazón, A.; Martinez-Mayorga, K. Bias-Inducing Allosteric Binding Site in Mu-Opioid Receptor Signaling. SN Appl. Sci. 2021, 3 (5), 566. 10.1007/s42452-021-04505-8.

(63) Sutcliffe, K. J.; Henderson, G.; Kelly, E.; Sessions, R. B. Drug Binding Poses Relate Structure with Efficacy in the μ Opioid Receptor. J. Mol. Biol. 2017, 429 (12), 1840–1851. 10.1016/j.jmb.2017.05.009.

(64) Trzaskowski, B.; Latek, D.; Yuan, S.; Ghoshdastider, U.; Debinski, A.; Filipek, S. Action of Molecular Switches in GPCRs - Theoretical and Experimental Studies. Curr. Med. Chem. 2012, 19 (8), 1090–1109. 10.2174/092986712799320556.

(65) Ricarte, A.; Dalton, J. A. R.; Giraldo, J. Structural Assessment of Agonist Efficacy in the μ-Opioid Receptor: Morphine and Fentanyl Elicit Different Activation Patterns. J. Chem. Inf. Model. 2021, 61 (3), 1251–1274. 10.1021/acs.jcim.0c00890.

(66) Ravindranathan, A.; Joslyn, G.; Robertson, M.; Schuckit, M. A.; Whistler, J. L.; White, R. L. Functional Characterization of Human Variants of the Mu-Opioid Receptor Gene. Proc. Natl. Acad. Sci. 2009, 106 (26), 10811–10816. 10.1073/pnas.0904509106.

(67) Zhou, Y.; Ramsey, S.; Provasi, D.; El Daibani, A.; Appourchaux, K.; Chakraborty, S.; Kapoor, A.; Che, T.; Majumdar, S.; Filizola, M. Predicted Mode of Binding to and Allosteric Modulation of the μ-Opioid Receptor by Kratom’s Alkaloids with Reported Antinociception *In Vivo*. Biochemistry 2021, 60 (18), 1420–1429. 10.1021/acs.biochem.0c00658.

(68) Schmid, C. L.; Kennedy, N. M.; Ross, N. C.; Lovell, K. M.; Yue, Z.; Morgenweck, J.; Cameron, M. D.; Bannister, T. D.; Bohn, L. M. Bias Factor and Therapeutic Window Correlate to Predict Safer Opioid Analgesics. Cell 2017, 171 (5), 1165–1175.e13. 10.1016/j.cell.2017.10.035.

(69) De Waal, P. W.; Shi, J.; You, E.; Wang, X.; Melcher, K.; Jiang, Y.; Xu, H. E.; Dickson, B. M. Molecular Mechanisms of Fentanyl Mediated β-Arrestin Biased Signaling. PLOS Comput. Biol. 2020, 16 (4), e1007394. 10.1371/journal.pcbi.1007394.

(70) Gillis, A.; Gondin, A. B.; Kliewer, A.; Sanchez, J.; Lim, H. D.; Alamein, C.; Manandhar, P.; Santiago, M.; Fritzwanker, S.; Schmiedel, F.; Katte, T. A.; Reekie, T.; Grimsey, N. L.; Kassiou, M.; Kellam, B.; Krasel, C.; Halls, M. L.; Connor, M.; Lane, J. R.; Schulz, S.; Christie, M. J.; Canals, M. Low Intrinsic Efficacy for G Protein Activation Can Explain the Improved Side Effect Profiles of New Opioid Agonists. Sci. Signal. 2020, 13 (625), eaaz3140. 10.1126/scisignal.aaz3140.

(71) Li, X.; Guo, Y.; Li, J.; Yu, Z.; Cheng, J.; Ren, F.; Jia, H.; Zhang, Y.; Cui, S.; Zhang, T.; Shi, W. Discovery and Structural Explorations of G-Protein Biased μ-Opioid Receptor Agonists. ChemMedChem 2022, 17 (24). 10.1002/cmdc.202200416.

(72) Gutridge, A. M.; Robins, M. T.; Cassell, R. J.; Uprety, R.; Mores, K. L.; Ko, M. J.; Pasternak, G. W.; Majumdar, S.; Van Rijn, R. M. G Protein-biased Kratom-alkaloids and Synthetic Carfentanil-amide Opioids as Potential Treatments for Alcohol Use Disorder. Br. J. Pharmacol. 2020, 177 (7), 1497–1513. 10.1111/bph.14913.

